# Intracellular mechanical fingerprint reveals cell type-specific mechanical differences

**DOI:** 10.1101/2023.08.10.552769

**Authors:** Till M. Muenker, Kerstin von Roden, Bart E. Vos, Timo Betz

## Abstract

Living cells are complex entities that perform various tasks with astonishing robustness. While the direct dependence of biological processes on controlled protein expression is well established, we have only begun to understand how intracellular mechanical characteristics guide and support biological function. This is in stark contrast to the expected functional role that intracellular mechanical properties should have for many core cellular functions such as organization, homeostasis, and transport. From a mechanical point of view, cells are complex viscoelastic materials that are continuously driven out of thermodynamic equilibrium, which makes both a physical measurement and mathematical modeling of their properties difficult. Here, we define a “mechanical fingerprint” that can not only characterize the intracellular mechanical state, but also carve out the mechanical differences between cell types with the potential to relate these to proper cell function. By analyzing the frequency-dependent viscoelastic properties and intracellular activity of cells using microrheology, we distilled the complex active mechanical state into just 6 parameters that comprise the mechanical fingerprint. The systematic investigation of the fingerprint illustrates a parameter tuning that could be related to functional cellular requirements. However, the full potential of the mechanical fingerprint is given by a statistical analysis of its parameters across all investigated cell types, which suggests that cells can modify mechanical parameters in a correlated way to position their intracellular mechanical properties within a well-defined phase-space that is spanned between activity, mechanical resistance, and fluidity. This paves the way for a systematic study of the interdependence of biological function and intracellular active mechanics.

## 1 Main

### 1.1 Introduction

Cells are highly complex, heterogeneous materials that tightly control their viscoelastic characteristics by precise tuning of the active mechanical properties to maintain a non-equilibrium mechanical steady state. This mechanical homeostasis requires a continuous consumption of metabolic energy, mostly in the form of ATP, and is maintained while the system undergoes various dynamic functions ranging from tissue morphogenesis and patterning^1, 2^ to organelle positioning,^3^ differentiation,^4^ migration,^5, 6^ and polarization.^7^

Although it is well-acknowledged that intracellular mechanical properties are key to understanding major cellular functions, most studies have measured cellular mechanics from outside of the cell using e.g. AFM,^8, 9^ micropipette aspiration,^10^ traction force microscopy,^11^ tether pulling,^12^ magnetic twisting cytometry,^13^ or deformability cytometry.^14^ While these studies allowed deep insights into the relevance of cell mechanics for supracellular biological processes by probing predominantly the stiff actin cortex that surrounds the cell, inferring information about the mechanical properties of the cytoplasm remained difficult.^15^ This experimental gap has recently been bridged by active intracellular rheology experiments that are based on optical as well as magnetic tweezers.^16–19^ For example, a recent study has shown that the cytoplasm softens during mitosis,^20^ which might be key for the proper functioning of this process.

While a large body of work has focused on the viscoelastic properties of cells, it has become increasingly clear that the well-controlled active force generation of living cells has to be integrated to obtain a full intracellular mechanical description. Recent studies have therefore focused on investigating these non-equilibrium, active properties of cells.^21–23^ These active, energy-consuming cellular processes are a key component to maintain the intracellular organization despite the ever-dispersing forces of entropy.^16, 24^ While intracellular transport continuously organizes membrane and organelle architecture via motor activity, the dynamic growth and disassembly of cytoskeletal filaments ensures global processes such as migration,^25^ proliferation,^26^ or wound healing.^27^ Further, recent studies have shown that dynamic waves of actin filaments also organize mitochondria position during cell division^28^ and that cytoplasmic forces can reorganize nuclear condensates,^29^ thus emphasizing the key importance of active forces for fundamental cellular function.

Despite the generally accepted relevance of intracellular active mechanics for the functioning of entire organisms, it remains difficult to quantify and even harder to model theoretically. From a physical perspective, cells behave as viscoelastic materials where timescales are key information to correctly identify the dominating mechanical principles. Therefore, a full quantitative description requires microrheological information, typically provided in the form of a frequency-dependent, complex shear modulus *G*\*(*f*).^30^ Although this quantification of the viscoelasticity is experimentally accessible, its interpretation requires theoretical models to become physically meaningful and to reduce the complexity of the data. However, while simple mechanical analogies, such as Kelvin-Voigt or Maxwell models lack the complexity to describe the observed behavior, more elaborate models, such as chains of springs and dashpots, require many parameters and thus do not reduce the complexity significantly. This has led to double power-law approaches, which are motivated by fractional Generalized Kelvin-Voigt (fGKV) models.^31^ In recent work, these have been successfully used to quantify the viscoelastic properties of fibroblasts,^32^ endothelial,^33^ and epithelial cells^20^ and they reduce the complex mechanical behavior to just four parameters. Recent efforts to include active force generation in a mechanical characterization were able to describe the active energy injected into the cellular system, also by a power-law^20^ that is defined by two additional parameters.

These insights raise the question to what extent living cells actively control their intracellular mechanical properties, or if these are simply byproducts of the cytoplasmic composition. Assuming a functional relevance, the parameters may vary considerably between cells and might even allow to identify different cell types and cellular situations in a fingerprint-like fashion. The relevance of this 6 parameter description was recently established by showing systematic changes during cell division, and to perform new statistical analysis of stochastic particle trajectories.^20, 34 –36^ In this study, we use optical tweezers-based active microrheology and passive observation of intracellular particle motion to demonstrate that cell types can be successfully distinguished in a pairwise comparison to position cells on 3D phasespace that reflects activity, rigidity, and fluidity.

### 1.2 Results and Discussion

#### Mechanical parameter set quantifies active mechanics

Intracellular viscoelastic material properties are commonly quantified by a frequency-dependent, complex shear modulus that we directly determine via optical tweezers-based active microrheology (M 1). Here, a custom-built optical tweezers setup (Figure 1a, M 2) exerts well-defined oscillatory forces onto phagocytosed 1 µm sized probe particles inside cervical cancer cells (HeLa). In this study, probe particles were added to the cell culture medium and taken up by the cells via endocytosis. Thus, they were surrounded by a membrane, and the obtained mechanical properties may serve as a proxy for membrane-bound organelles of similar size. To determine the resulting bead displacement, a stationary, weak detection laser monitors the particle position with nanometer precision. Applying sinusoidal forces (Figure 1b) at increasing frequencies of 1, 2, 4, 8, 16, 32, 64, 128, 256, 512, and 1024 Hz, while maintaining a constant trapping-laser oscillation amplitude of 200 nm, allows determination of the mechanical response function *χ*(*f*). This response function is then used to calculate the complex shear modulus according to *G*\*(*f*) = 1/(6*πRχ*(*f*)), where *R* denotes the radius of the probe particle (Figure 1c). *G*\*(*f*) is a complex quantity where the real part *G*′(*f*) (storage modulus) describes the elastic properties of the cell and the imaginary part *G*″(*f*) (loss modulus) describes the viscous properties (Figure 1c).

**Figure 1:**
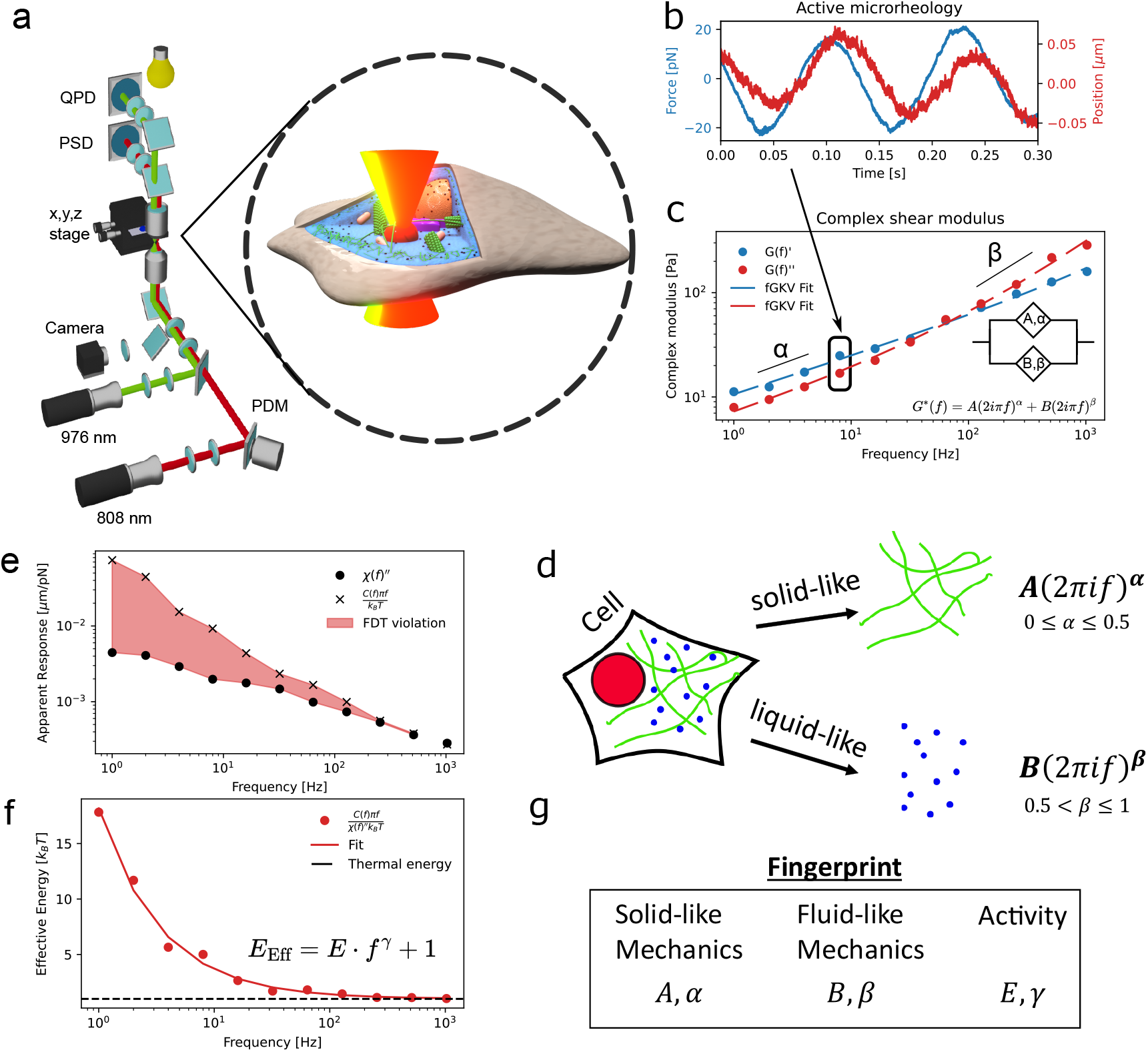
a) Viscoelastic properties and activity are measured within the cytoplasm of cells using a custom-built optical tweezers setup. b) Sinusoidal forces (blue) at varying frequencies are applied to a probe particle while the particle displacement (red) in response to this force is monitored. c) Repeating this procedure at different frequencies allows to calculate viscoelastic material properties in terms of the complex shear modulus *G*\*(*f*) which consists of the storage modulus (blue) and a loss modulus (red). Both storage and loss modulus can collectively be fitted using a generalized fractional Kelvin-Voigt (fGKV) model (inset). d) This fGKV model consists of two power-laws, which in a simplified way can be pictured as a more solid-like material class resembling polymeric filaments and a more fluid-like material class (crowded molecules). e) Using an additional passive measurement, violation of the fluctuation-dissipation theorem can be directly visualized. (red area) f) The effective energy quantifies intracellular activity and can be fitted with a two-parameter power-law. g) Complex and frequency-dependent intracellular active mechanical properties can be reduced to a fingerprint of just 6 parameters, which describe the intracellular mechanical state.

To model the shear modulus, we follow the naive hypothesis that the cytoplasm consists of two main material classes, namely an elastic solid-like material that is generated by a sparse, connected polymer network, and a more liquid-like material that fills the bulk space of the cytoplasm (Figure 1d). Generally, such a combination of two complex materials can be approached by a linear combination of two springpots that leads to a double power-law model consistent with classical rheology data^20, 31 –33^ :

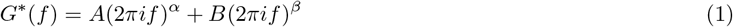

This complex function can then be used to simultaneously fit both the real and imaginary parts of the experimentally obtained shear modulus to determine the prefactors and power-law exponents. As shown in Figure 1c the model fits the experimental data well. *R*^2^-values of the fits are given for raw data (Appendix Table 16) and bootstrapped data (Appendix Table 15). Consistent with the initial hypothesis, we identify a region with a power-law exponent between 0 and 0.5 that we attribute to a more solid-like material (ideal solid has exponent 0), and a second power-law that is between 0.5 and 1 (ideal fluid has exponent 1) corresponding to a liquid-like material (Figure 1, right).

In addition to intracellular mechanics, the generated active forces are another key determinant of the intracellular state. As cells consume metabolic energy to drive motor proteins and many other cellular functions, cells are far from thermodynamic equilibrium. Experimentally, this is directly reflected in the motion of tracer particles within the cell. In thermodynamic equilibrium, the random motion of particles depends only on material properties and temperature, which is manifested by the fluctuation-dissipation theorem:^37^

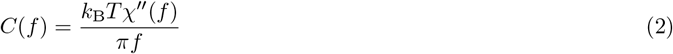

Here, *C*(*f*) is the power spectral density, quantifying the spontaneous fluctuations of the particle, *χ*(*f*) is the mechanical response function, and *k*_B_*T* is the thermal energy (M 1.2). In active, living cells, particle mobility is beyond motion explainable by thermal fluctuations^38^ (Figure 1e), as cytoskeletal rearrangement and transport processes also contribute to the motion of probe particles. An elegant way to summarize all active and thermal forces is by introducing an effective energy *E*_Eff_ ^37^(M 1). Intuitively, this effective energy reflects the ratio between observed particle motion *C*(*f*) and thermodynamically predicted motion that reflects the dissipative material properties as quantified by *χ*^″^(*f*):

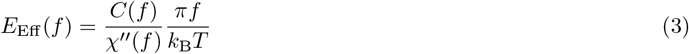

It should be noted that this idea of an effective energy implies that the active forces generated by the cell share statistical characteristics with thermal forces. In this sense, we use the effective energy rather as a phenomenological quantity that measures the extent of non-equilibrium mechanical driving of the system. The free particle fluctuation spectrum *C*(*f*) is acquired by turning off the trapping laser while using the lower-power detection laser to monitor the particle position. The resulting effective energy measured for a representative HeLa cell (Figure 1f) shows an increased influence of active forces in the low-frequency regime, while the effective energy converges to the thermal limit of 1 *k*_B_*T* for higher frequencies. This directly demonstrates that metabolic energy dominates particle motion at long timescales, while assuming thermodynamic equilibrium at shorter timescales remains a valid approximation. This asymptotic behavior at short time scales is to be expected as motor proteins operate at longer time scales,^39^ and has previously been shown experimentally.^16, 40 –42^ This behavior was also used for an additional calibration method during optical tweezers experiments as described in Appendix 1.1. Interestingly, this suggests that the question about non-equilibrium processes can be reduced to a question of timescales, which is a potential explanation why this subject is so controversially discussed in literature.

Similar to the mechanical properties, the effective energy can also be fit by a power-law approach (Figure 1f):

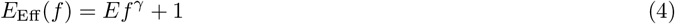

This allowed us to describe the active forces generated by metabolic processes within the cell using only two parameters. Together with the four parameters describing the viscoelastic response, these quantities define a set of six parameters that characterize the intracellular mechanical state of a cell, as shown for the active mechanical properties of the cytoplasm in HeLa cells in Figure 1f. As we show in the following, the set of these 6 parameters can be interpreted as a mechanical fingerprint of a cell (Figure 1g).

In this study, we explore the variation of the mean active mechanical quantities *G*′, *G*″, and *E*_Eff_ across different cell types or conditions. To obtain a robust and reliable distribution of these curves, we apply a bootstrapping scheme to the recorded single-cell data (see M 3.1). The bootstrapped data is then subjected to respective model fitting, providing estimations for the distributions of the mean fingerprint parameters for each condition. Unless otherwise specified, all presented data is based on the bootstrapped data.

#### 1.2.1 The cytoskeleton selectively affects the mechanical fingerprint

After establishing that the mechanical fingerprint provides an adequate description of the intracellular mechanical state of HeLa cells, we investigated whether individual fingerprint parameters can be selectively modulated by pharmacological perturbation of the cytoskeleton (Figure 2a). Cellular mechanics are commonly attributed to the actin cytoskeleton, which forms the cell cortex and a variety of intracellular structures. To assess the contribution of actin, we treated HeLa cells with 10 *µ*g*/*mL Cytochalasin B (CytoB or CB) and 1 *µ*M Latrunculin A (LatA), two compounds that interfere with actin polymerization through distinct mechanisms (Figure 2a).

**Figure 2:**
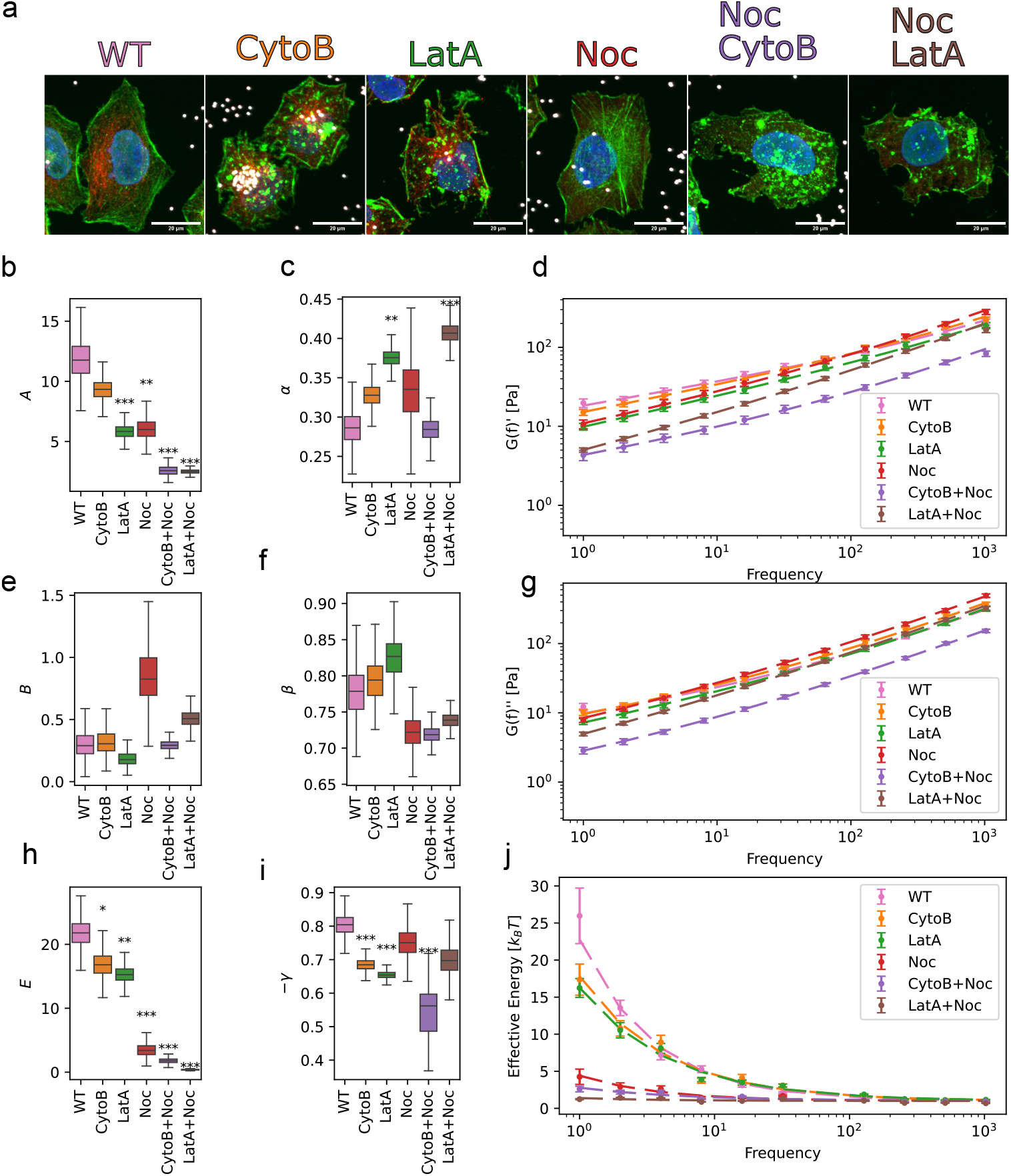
Effect of cytoskeletal perturbations on the intracellular mechanical fingerprint of HeLa cells. a) Composite images of HeLa WT cells treated with Cytochalasin B (CytoB), Latrunculin A (LatA), Nocodazole (Noc), Cytochalasin B + Nocodazole (CytoB+Noc), and Latrunculin A + Nocodazole (LatA+Noc). DNA stained with Hoechst in blue, microtubules stained with *α*-tubulin antibody in red, actin stained with phalloidin in green, and probe particles in white. Scale bar: 20 µm b-j) Changes in the fingerprint parameters and corresponding rheological quantities following cytoskeletal perturbation. The dashed lines in plots for *G*′, *G*″ and *E*_Eff_ show the models depicted in Figure 1c,f using the mean fit parameters *A, B, E, α, β* and *γ*.

CytoB treatment resulted in a modest but statistically non-significant decrease of the solid-like prefactor *A* (*p* = 0.08; Figure 2b), suggesting only a minor reduction in intracellular elasticity. Consistently, both the storage and loss moduli were slightly reduced, particularly at low frequencies (Figure 2d,g). LatA treatment produced a qualitatively similar but substantially stronger response. The solid-like prefactor *A* was significantly reduced (Figure 2b), while the power-law exponent *α* increased (Figure 2c), indicating a softer and more fluid-like intracellular environment. These observations are qualitatively consistent with previous AFM studies that reported a reduction in cortical stiffness following actin depolymerization.^43, 44^ In addition, both actin-disrupting treatments significantly reduced intracellular activity, as reflected by a decrease in the effective-energy prefactor *E* (Figure 2h,j). The more pronounced effect of LatA is consistent with its mechanism of action. While CytoB primarily caps filament ends and therefore leaves stabilized actin filaments largely intact, LatA directly binds to actin monomers, resulting in more extensive depolymerization of the actin network.^45^ Consequently, CytoB substantially reduced intracellular activity while producing only a modest reduction in intracellular mechanical resistance, whereas LatA strongly affected both the mechanical and active components of the fingerprint. This interpretation is supported by immunostaining experiments (Figure 2a and Appendix Figure 2), which revealed extensive reorganization of the actin cytoskeleton following both treatments. In both cases, filamentous actin structures became disrupted and the actin signal appeared redistributed into clustered structures. Compared with CytoB, LatA produced a more complete loss of recognizable filamentous actin, consistent with its stronger mechanical effect. As intracellular activity remained well above thermal levels after actin disruption, we next investigated the contribution of microtubule-associated processes. Since the probe particles are internalized by endocytosis and are typically enclosed within late endosomes or lysosomes, they are expected to interact strongly with microtubule-based transport processes. We therefore treated HeLa cells with 10 *µ*g*/*mL Nocodazole to depolymerize the microtubule cytoskeleton. As expected, immunostainings for *α*-tubulin revealed a strong reduction of filamentous microtubule signal (Figure 2a, Appendix Figure 2). Nocodazole treatment caused a strong reduction of the solid-like prefactor A (Figure 2b) and overall reduction of storage and loss modulus (Figure 2d,g), and dramatically decreased intracellular activity from *E* = 23*k*_*B*_*T* to *E* = 4*k*_*B*_*T* (Figure 2h,j). Interestingly, this behavior differs from previous AFM studies probing cortical mechanics, which reported either no significant change in stiffness upon microtubule depolymerization^46^ or even a stiffening response.^44^ This discrepancy likely reflects the different mechanical compartments probed by the two methods. Whereas AFM predominantly probes the actin-rich cell cortex, our measurements characterize the mechanical environment experienced by membrane-bound intracellular particles, which are expected to interact strongly with microtubule-based transport processes. Consistent with the pharmacological treatment, immunostaining confirmed the near-complete loss of the microtubule network following Nocodazole treatment, while revealing a qualitative increase in actin stress fibers (Figure 2a, Appendix Figure 2). Together, these findings support the interpretation that microtubule-associated processes make a major contribution to the active mechanical properties experienced by phagocytosed particles.

To this point, the data suggest that both filament types, actin and microtubules, contribute to the overall cytoplasmic mechanical properties, even though with different proportions. Yet, no treatment by itself was able to fully suppress intracellular activity. To test whether additional processes could contribute, we performed double disruption of actin and microtubules using the combined treatment with CytoB + Noc and LatA + Noc. Both conditions led to a reduction of cytoplasmic microtubules as well as dissolution of actin fibers and actin aggregation (Figure 2a and Appendix Figure 2). In both cases, the solid-like properties decreased beyond the levels previously observed by single disruptions (Figure 2b,d). The same can now also be observed for intracellular activity as quantified by E, which is reduced to about 2*k*_*B*_*T* for CytoB + Noc and down to almost thermal energy 1*k*_*B*_*T* for LatA + Noc treatment (Figure 2 h,i,j). The combined disruption of actin and microtubules is sufficient to suppress most of the intracellular activity experienced by phagocytosed particles.

These results demonstrate that the mechanical fingerprint can be used to dissect the effect of cytoskeletal drugs on different properties. At this point, it is important to stress that these experiments were performed on phagocytosed beads, likely engulfed in a membrane, which is an adequate proxy for membrane-bound organelles. The observed relations here can not be readily translated to inert particles. Other studies have found, for example, that the mechanical properties felt by inert probe particles are largely influenced by molecular crowding. Here, an increased intracellular activity led to a decreased overall stiffness.^47, 48^

#### 1.2.2 Separating cell types by their mechanical fingerprint

The strong sensitivity of the solid-like prefactor *A* and the effective-energy prefactor *E* to cytoskeletal perturbations suggests that these parameters may be subject to cell-type-specific regulation. We therefore investigated whether cells with distinct biological functions and mechanical demands exhibit different mechanical fingerprints by comparing HeLa cells with two additional cell types (Figure 3a). First, we hypothesized that a muscle cell needs to have an increased intracellular mechanical stiffness for its function as a mechanical force generator. This suggests that muscle cells will be tuned to differ in the solid-like material properties with a higher prefactor A.

**Figure 3:**
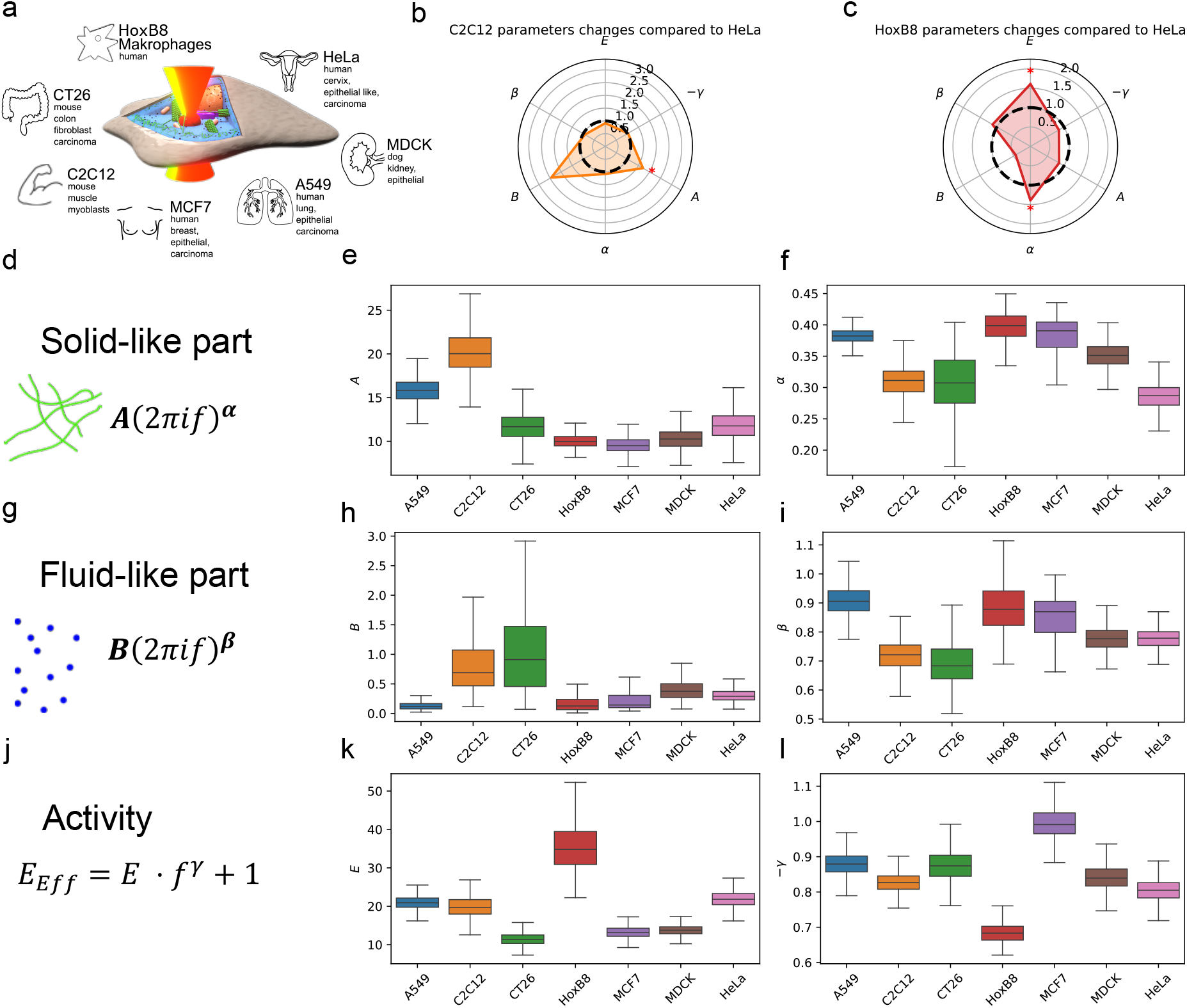
a) The mechanical fingerprint of 7 different cell types was determined by measuring their complex shear modulus and effective energy. b) Comparing HeLa cells to C2C12 muscle cells shows that muscle cells are overall stiffer (higher *A*). c) The comparison between HeLa cells and macrophages shows that macrophages have a higher intracellular activity (increased *E*) but are also more liquid-like and softer (increased *α, β* and decreased *B*) d-l). Using the fingerprint, active mechanical differences between cell types can be compared according to changes in the more solid-like material properties, the more liquid-like material properties, and intracellular activity.

As seen in Figure 3b, this is exactly what we found when comparing the cervical cancer HeLa cells with a murine-derived myoblast cell line (C2C12). The solid-like material stiffness, as quantified by *A*, significantly increased from about 13 to 20 (Figure 3b,d,e). Additionally, the fluid-like material prefactor *B* increased, indicating that the fluid-like material class also has more resistive properties (Figure 3b,g,h).

Next, we hypothesized that immune cells would have increased active forces in the cytoplasm and render it more liquid-like to be able to move more quickly through small constrictions as found in the extracellular space. This hypothesis was tested using an immune cell model. Macrophages were obtained from a hematopoietic progenitor cell line that expresses HoxB8 under an estrogen promoter.^11, 13^ Upon removal of estrogen, the cells differentiate within 3 days to macrophages that we term HoxB8 cells in the following for convenience. Again, our experimental analysis of the mechanical fingerprint is consistent with the prediction based on the biological function of macrophages (Figure 3c). Firstly, the effective energy prefactor *E* increased, indicating a raised intracellular activity. Secondly, the solid-like exponent *α* significantly increased, while the viscosity-defining prefactor *B* of the liquid-like material property collapsed (Figure 3f,h,k), indicating that the solid-like material class behaves more liquid-like, allowing the cytoplasm to quickly dissipate any large-scale deformation of the whole cell. Our results are in good agreement with measurements obtained previously on the whole cell level.^49^

These results suggest that differences in cellular function are associated with differences in mechanical fingerprint parameters. Muscle cells, requiring higher mechanical resistance, show a large solid-like prefactor, while immune cells that need to migrate quickly and deform easily are more liquid-like with lower viscous dissipation and generate a larger cytoplasmic active energy.

#### 1.2.3 The mechanical fingerprint is different across cell types and species

Motivated by the finding that different cell types have distinct mechanical properties as quantified by the mechanical fingerprint, we wondered whether this approach could be used as a strategy to differentiate between cell types in terms of mechanical properties. In detail, we ask if at least one of the 6 parameters of the mechanical fingerprint is statistically different when comparing different cell types. Hence, we decided to determine the 6-parameter fingerprint for 7 different cell types (Figure 3a), of various functions and from various species. In addition to the already introduced human HeLa, murine C2C12 and HoxB8 cells, we measured the fingerprint of human lung epithelial cells (A549), a murine fibroblast-like colon carcinoma cell line (CT26), a canine kidney epithelial cell line (MDCK), and a non-invasive human breast cancer cell line (MCF7). Figure 3d-l summarizes the six fingerprint parameters across all seven investigated cell types. The observed trends are broadly consistent with previous studies of cell mechanics. For example, AFM measurements probing cortical stiffness reported a similar ordering of A549, MDCK, and MCF7 cells (A549: 1.6 kPa *>*MDCK: 0.56 kPa *>*MCF7: 0.25 kPa).^50^ As expected, the absolute stiffness of the cortex is much higher than that found in the cytoplasm. Notably, the same ordering is observed for the solid-like prefactor *A* (Figure 3e), despite the fact that our measurements probe the mechanical environment experienced by intracellular tracer particles rather than the cell cortex directly.

In a first analysis, we looked at all possible pair-wise combinations of cells to test our hypothesis that the 6 parameters are sufficient to differentiate between cell types. Indeed, in most of the 21 possible combinations we found at least one of the fingerprint parameters to be different (Figure 4a, Appendix 2.3). However, this analysis does not take into account the extent of the significant difference. To approach this question more systematically, we use a variant of the z-score, where we normalize the difference in the mean values by twice the geometric average of the standard deviation of the bootstrapped data. The z-score, hence, quantifies the extent of deviation between cells that is carried by a single parameter. For example, the z-score of A between HeLa and CT26 is:

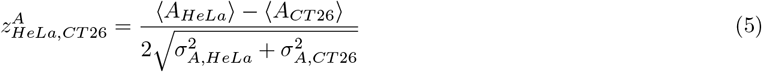

**Figure 4:**
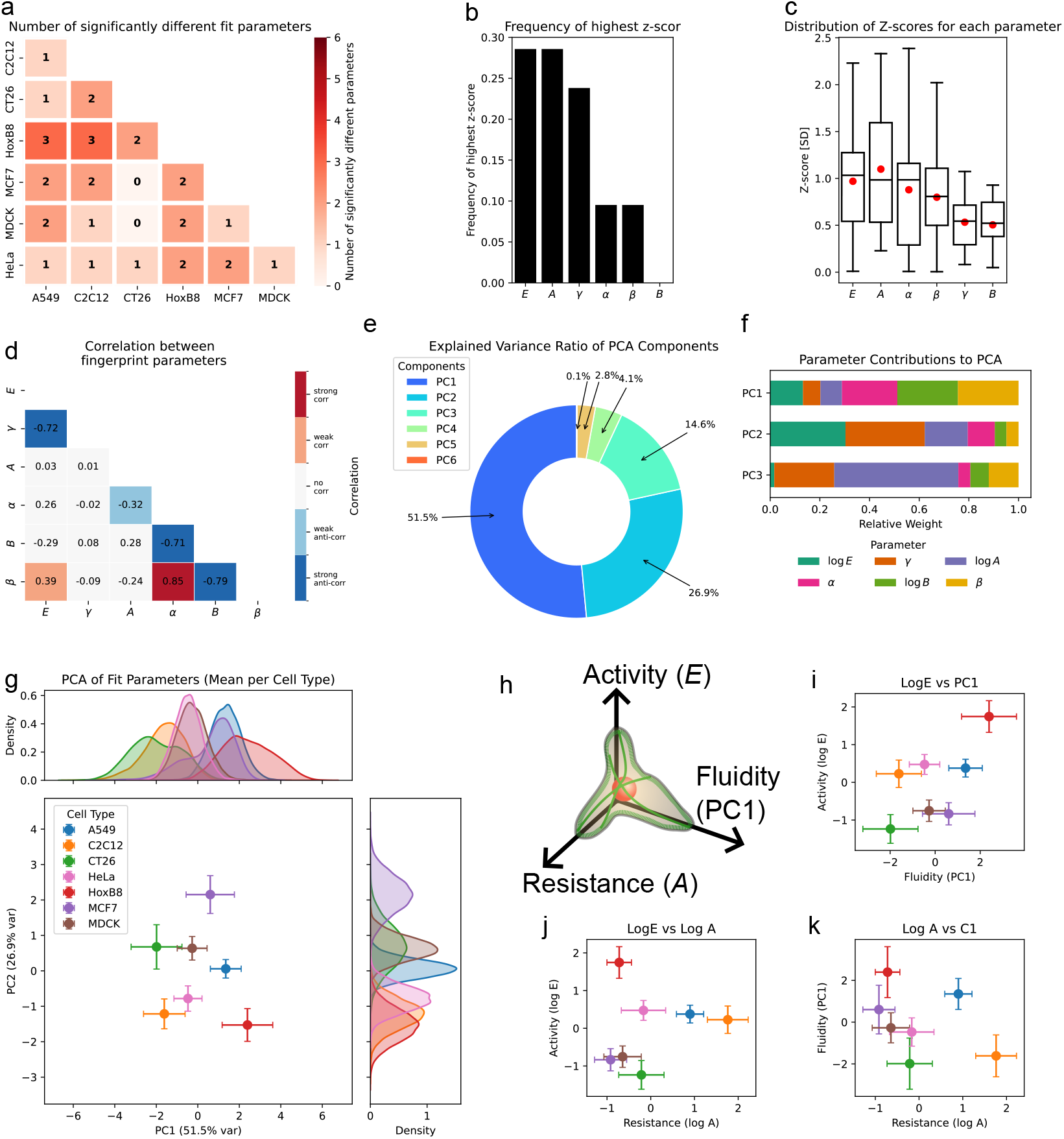
a) Number of significantly different fingerprint parameters in pairwise cell type comparisons. b) Frequency of parameters showing the highest z-score. c) Distribution of z-score for all parameters for pairwise cell comparison. d) Correlation analysis between fingerprint parameters shows that not all parameters are varied independently. e) Explained variance ratio of the different principal components (PC) of the principal component analysis (PCA). f) Relative contribution of the fingerprint parameter to the first three principal components. PC1 mainly consists of *α, β* and log *B*, PC2 mainly consists of log *E* and *γ* and PC3 to 50% of log *A*. g) Plotting all cell types according to the first two principal components PC1 and PC2 shows that two parameters are already sufficient to distinguish between most cell types. h,i,j,k) Qualitative phase diagram of the active mechanical space. Activity, mainly captured by parameter *E*, resistance, dominated by *A*, and solid-liquid switching can be described as fluidity that is determined by principal component 1 (PC1) are varied among different cell types. Using this three-dimensional space allows identification of physical differences among different cell types.

Here, any value greater than 1 can be considered to mark a significant difference, and the larger the z-score, the better the two cells can be separated according to the respective parameter. We then calculated the z-score for all 6 fingerprint parameters across all 21 different pairs of cells. In a first step, we studied which of the parameters was best to differentiate between the cells. Figure 4b shows how often each parameter was the best discriminator. We see that *E* and *A* are the most important parameters to distinguish between different cell types as they together show the highest z-score in about 55% of the cases. As they mainly characterize the solid-like material properties and overall activity, this result implies that these properties are highly dependent on cell type and function. Additionally, we directly plot the z-scores of all cell pairs for the different parameters (Figure 4c). In contrast to the previous quantity, here the value of the z-scores becomes important. Despite *E* and *A* having most frequently the highest z-score, we also observed that the average z-score for all parameters was similar or smaller than 1, suggesting that the combination of all 6 parameters is relevant for characterizing the mechanical state of cells. The similarity in average z-scores may also indicate that there are correlations between these parameters, which could be due to underlying biological mechanisms.

To further explore this possibility, we looked at the correlation values (c.v.) (M 3.4) between the fingerprint parameters across all measurements as depicted in Figure 4d. We define an absolute c.v. below 0.3 as no (anti)correlation, between 0.3 and 0.6 as weak (anti)correlation, and above 0.6 as strong (anti)correlation. While for most parameter pairs no or only weak correlations are observed, we find a strong dependency between parameters *B, β* and *α*, as well as between *E* and *γ*. While *α* and *β* are positively correlated (0.85), both are negatively correlated with *B* (−0.71 and −0.79). A simultaneous variation of *α, β*, and *B* might be a way for the cell to switch between a solid- and fluid-like state. This finding also implies that, even though the two-power-law approach is able to perfectly describe the observed viscoelastic properties, a more simple model might also be able to fit the data, as some parameters are not fully independent. In this simpler model, the cytoplasm does might not consist of two independent classes of materials, but instead a deeper underlying mechanism might regulate the low-frequency elastic-like behavior just as the high-frequency more fluid-like behavior.

To see if we can indeed further reduce the number of parameters required to describe the active intracellular mechanical state, we performed a principal component analysis (PCA) with all three exponents *α, β*, and *γ*, and the logarithm of prefactors log *A*, log *B* and log *E* (M 3.5). The explained variance ratio of each component is presented in Figure 4e. Already 93% of the variance can be explained using only the first 3 components, and 78.4% can even be explained with just 2 components. This supports the hypothesis that few underlying biological mechanisms act as “hyperparameter”, which tune the mechanical fingerprint of the intracellular space.

Each principal component is a superposition of the normalized fingerprint parameters. We therefore wondered which fingerprint parameters dominate the first 3 principal components. Looking at the relative contributions (Figure 4f, Appendix 2.1), we find that the first component PC1, explaining 51.5% of all variance in data, mainly consists of fingerprint parameters *α, B*, and *β* (22%, 24%, 24%). Finding these parameters combined in a single principal component is expected, as they also show strong correlation (Figure 4d), but strikingly, even though neither parameter by itself appeared to be dominant in characterizing the intracellular state in the previous analysis (Figure 4b,c), the combined component PC1 seems to be the most relevant principal component. The second component PC2 consists mainly of the fingerprint parameters related to the intracellular activity *E* and *γ* (30.3% and 32%), and the third one largely consists of parameter *A* (50%), characterizing the overall resistance of the cytoplasm felt by phagocytosed particles.

To determine whether PC1 and PC2 are already sufficient to distinguish between cell types, we plot the cell types according to these two parameters (Figure 4g). We observe that most cell types appear to separate from one another, indicating that PC1 and PC2 are effective in distinguishing between them.

As the first three principal components are sufficient to distinguish most of the investigated cell types, we sought to assign a physical interpretation to them. The first principal component, PC1, is dominated by the parameters *α, β*, and *B*. Because these parameters are strongly correlated across the investigated cell types, an increase in PC1 corresponds to a transition from a more elastic, solid-like response toward a more dissipative and fluid-like mechanical state. We therefore interpret PC1 as a measure of cellular fluidity. In contrast, the second principal component is dominated by the intracellular activity parameter *E* and *γ*. As the overall amplitude of effective energy is mainly influenced by a variation in *E* and less by *γ*, an increase in PC2 reflects an increase in intracellular activity. Lastly, as the third component is mainly represented by parameter *A*, which in turn is typically varied over an order of magnitude higher values than *B*, a change of this component can be attributed to an overall change of absolute complex modulus and thus a change of overall resistance.

To avoid this ambiguity of having principal components consisting of multiple parameters while preserving the main features revealed by the principal component analysis, we propose a qualitative three-dimensional phase space spanned by intracellular activity (primarily represented by log *E*), cellular resistance (primarily represented by log *A*), and fluidity (represented by PC1) (Figure 4h).

Mapping the different cell types into this phase space reveals distinct mechanical phenotypes (Figure 4i,j,k). HoxB8 macrophages exhibit the highest intracellular activity while simultaneously occupying the most fluid-like region of the phase space and, together with MCF7 cells, displaying comparatively low resistance. In contrast, C2C12 myoblasts are characterized by high resistance and one of the most solid-like mechanical states. These observations suggest that different cell types occupy distinct regions of the active-mechanical phase space, potentially associated with their physiological functions.

Having established that different cell types can be distinguished by their active mechanical state, an important next question is how these mechanical properties relate to cellular function. In particular, it would be interesting to determine whether parameters such as migration speed, contractility, or proliferation correlate with specific regions of the phase space. However, quantitative functional data are highly dependent on experimental conditions, including culture medium, substrate properties, and assay design, making direct comparisons across the investigated cell types difficult. As one example, wound-healing experiments have reported higher migration speeds for HeLa cells than for MCF7 cells,^51^ which is qualitatively consistent with the higher intracellular activity observed for HeLa cells in our measurements. To obtain a first indication of possible structure-function relationships, we grouped the investigated cell types according to species, epithelial or mesenchymal character, cancer status, metastatic potential, and presumed migratory potential (Appendix Figure 3 Appendix Table 17). However, no clear correlations emerged from this analysis, emphasizing the need for future studies that combine intracellular rheological measurements with quantitative functional assays.

### 1.3 Conclusion

In this study, we introduced a mechanical fingerprint that quantitatively describes the active mechanical state of the cytoplasm using only six parameters. By combining active and passive intracellular microrheology with a minimal physical description, we reduced the complex frequency-dependent viscoelastic and active properties of living cells to a compact parameter set that captures intracellular mechanics across a broad range of timescales.

Using pharmacological perturbations of the cytoskeleton, we demonstrated that individual fingerprint parameters reflect distinct physical aspects of the intracellular environment. While disruption of actin filaments, especially using Latrunculin A, primarily affected the solid-like mechanical response, microtubule depolymerization strongly reduced intracellular activity. Combined disruption of both filament systems reduced active fluctuations to nearly thermal levels, illustrating that the fingerprint can disentangle different contributions to intracellular active mechanics. Comparison with previous AFM studies revealed that actin depolymerization induces qualitatively similar softening responses at both the cortical and intracellular levels. In contrast, microtubule depolymerization produced markedly different effects, highlighting that intracellular microrheology using phagocytosed beads probes a mechanical compartment distinct from the actin-rich cell cortex and is particularly sensitive to microtubule-associated intracellular processes.

Extending the analysis across multiple cell types revealed that the mechanical fingerprint is not randomly distributed but instead exhibits characteristic differences between cell types. Muscle cells exhibited increased mechanical resistance and more solid-like properties, whereas macrophages displayed enhanced activity and increased fluidity. More generally, the statistical analysis revealed that the fingerprint parameters do not vary independently across the investigated cell types. Instead, changes in individual parameters are accompanied by systematic changes in others, suggesting that intracellular mechanical states occupy a constrained region of parameter space.

This coordinated tuning is reflected in a low-dimensional active mechanical phase space. Principal component analysis identified fluidity as a dominant mode of variation, while activity and mechanical resistance emerged as additional key descriptors. Together, these quantities define a three-dimensional framework that allows different cell types to be positioned according to their intracellular mechanical state. We propose that this phase space provides a useful basis for relating intracellular mechanics to biological function.

Several limitations should be considered when interpreting the presented results. The measurements were performed on phagocytosed probe particles that are likely enclosed within membrane-bound compartments and therefore probe the mechanical environment experienced by endosomes or lysosomes rather than the bulk cytoplasm directly. Consequently, the reported fingerprint reflects the active mechanics of membrane-associated intracellular structures and may differ from measurements obtained using inert tracer particles. Here, the mechanical properties are to a large extent influenced by the actin and microtubule cytoskeleton, while microtubule-associated processes emerge as major contributors to the measured fingerprint. Besides cytoskeletal organization, metabolic state, molecular crowding, and intracellular transport dynamics are likely to contribute to the observed fingerprint. Other studies probing inert particles in the cytoplasm have shown that intracellular mechanics can be strongly affected by molecular crowding.^47, 48^ Future studies combining intracellular rheology with metabolic perturbations or direct measurements of cellular energy metabolism will be required to disentangle the relative contributions of these factors.

Taken together, our results establish the mechanical fingerprint as a quantitative framework for characterizing intra-cellular active mechanics. We anticipate that this approach will enable systematic comparisons between cell types and physiological states and provide new opportunities to investigate how intracellular mechanics relate to cellular function and disease.

## 2 Material and Methods

### M 1 Active and passive microrheology in cells

In order to conduct active and passive microrheology measurements, we utilized the previously described optical tweezers setup (see Figure 1a). Cells were prepared following the protocols outlined in Materials and Methods M 4. Throughout the experiments, the cells were maintained at 37 ^°^C and 5% CO_2_ to ensure physiological conditions. We investigated a minimum of N=58 cells for each cell type or condition (see Appendix 2.2) over the course of at least n=3 consecutive days. For each cell, one particle was investigated. Data points were recorded at a sampling rate of 65 536 Hz.

#### M 1.1 Active microrheology

In active microrheology, the 808 nm trapping laser is employed to oscillate with a variable driving frequency *f*_*D*_, exerting a sinusoidal force *F* (*t*) onto a probe particle located inside a cell. Simultaneously, a second position detection laser records the position *x*(*t*) of the particle. The relationship between the particle’s position and the applied force is determined by the response function *χ*(*t*), as given by:

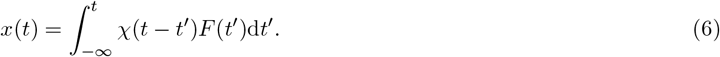

In the frequency domain, this convolution can be evaluated as a product, providing access to the response function:

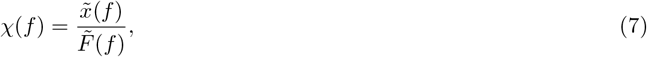

which characterizes the viscoelastic properties of the probed region. Here, 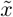 and 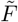 denote the Fourier transform of position and force.

To obtain the frequency-dependent response function, the probe particle is successively driven at different driving frequencies (*f*_*D*_ ∈ 1Hz, 2Hz, 4Hz, 8Hz, 16Hz, 32Hz, 64Hz, 128Hz, 256Hz, 512Hz, 1024Hz). At each frequency, a minimum of 3 periods was recorded. For frequencies above 2Hz, we recorded a maximum duration of 1 second. Utilizing the generalized Stokes-Einstein relation:^52^

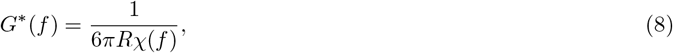

where *R* represents the particle radius, we determine the complex shear modulus *G*\*(*f*).

#### M 1.2 Passive microrheology

In passive micro rheology measurements, free particle fluctuations inside the cells are recorded without application of external forces. In this study, we employ the weak detection laser and back focal plane interferometry to track the bead displacement with high spatial and temporal precision. The particle position *x*(*t*) is tracked 3 consecutive times over a time span of 10 s for each cell investigated. For each time series *x*(*t*), we calculate the power spectral density (PSD)

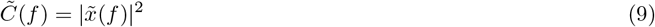

and average it over the 3 repetitions.

In equilibrium systems, the PSD is linked to the mechanical properties of the system via a generalized fluctuation-dissipation theorem (FDT):^52, 53^

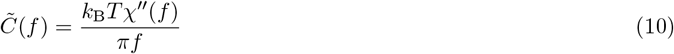

Here *k*_*B*_*T* is the thermal energy of the system and *χ*^″^(*f*) the imaginary part of the response function. In passive systems, this equation is valid and 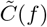 can be used to determine the mechanical properties of the system. Living cells, however, are non-equilibrium systems. Metabolic energy leads to stronger particle fluctuations 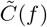 than one would expect for a passive systems. We define an effective energy by adding an active cellular energy *E*_*active*_ to the thermal energy:

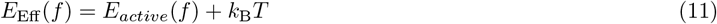

This is the energy required to explain the particle fluctuations 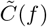 given the mechanical properties *χ*^″^(*f*) measured in active microrheology. This yields a new adapted version of the FDT:

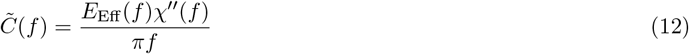

By separately measuring *χ*^″^(*f*) in active microrheology and 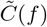 in passive microrhelogy, we gain access to the effective energy of the system and thereby the level of non-equilibrium:

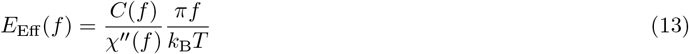

The effective energy quantifies the level of activity in the system. Any deviation from 1 is indicative for non-equilibrium.

### M 2 Optical setup

In this study, we use an optical tweezers setup that has been previously described^34^ to perform active and passive microrheology measurements and apply well-defined forces onto phagocytozed probe particles in living cells. Briefly, the setup is based on a home-built bright-field microscope equipped with a 60x NA 1.2 Objective, where we employ two different lasers. A 808 nm laser (LU0808M250, 808 nm, 250 mW Lumics GmbH, Berlin,331 Germany) is operated at moderate power (70 mW at sample plane) to allow for stable optical trapping and force application to probe particles at the focal plane. The lateral position of this laser can be precisely controlled with a piezo-based tilting mirror. Using a high NA=1.4 condenser, most of the transmitted laser light is collected. By projecting the back focal plane of the condenser onto a position sensitive detector (PSD), the trapping forces can be directly quantified.^54^ A second, stationary, laser (L976-PAG500, 976 nm, 500 mW, Thorlabs, New Jersey, USA) is operated at low laser power (0.3 mW at sample plane) to not apply any relevant trapping forces. This laser is utilized to monitor the bead displacement via back focal plane interferometry^55^ with nm precision. A schematic of this setup is depicted in Figure 1a.

### M 3 Statistical analysis

#### M 3.1 Bootstrapping

Bootstrapping is employed to get an estimate for the distribution of the mean cellular active mechanical properties. The following scheme is used:

- We have *i* experimental conditions, either different cell-types or drug treatments. For each condition *i*, we inves-tigate *N*_*i*_ different cells. For each cell, we obtain curves for *G*′,*G*″ and *E*_Eff_
- For each condition *i*, the bootstrapping data set is generated. *N*_*i*_ samples are drawn with replacement with consecutive mean calculations. This process is repeated *N*_bootstrap_=10.000 times for each quantity of interest.
- The resulting distribution of curves captures the distribution of the means.
- Each curve is subsequently fit with the respective model, which in turn also yields the distribution of the mean fit parameters.

#### M 3.2 Significance tests

Statistical comparisons between fit parameters obtained from the bootstrapping scheme (Section M 3.1) were performed as follows;^56^ a schematic is provided in Figure 1.

For each pairwise comparison of two groups *G*_1_ and *G*_2_, we exploit that bootstrapping yields a full sampling distribution of each fit parameter rather than a single point estimate. Let 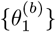 and 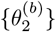 denote the bootstrap distributions of a given parameter for the two groups. We form the difference distribution

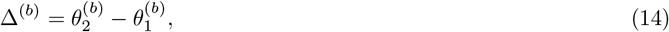

and estimate the two-sided *p*-value as

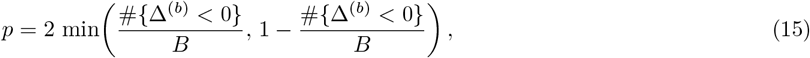

where *B* is the number of bootstrap replicates. Intuitively, this tests whether zero is contained in the bulk of the difference distribution. If the two groups differ substantially, the entire distribution is shifted away from zero, and the *p*-value is small.

Because multiple pairwise comparisons are performed simultaneously for each parameter, we control the familywise error rate using the Holm–Bonferroni sequential correction.^57^ Significance levels are then assigned to the adjusted *p*-values as: *p* ≤ 0.001 (*∗*), *p* ≤ 0.01 (**), *p* ≤ 0.05 (*), and *p* > 0.05 (n.s.).

#### M 3.3 Z-score

We use an adapted z-score to quantify how different a certain parameter is between two different cell types.

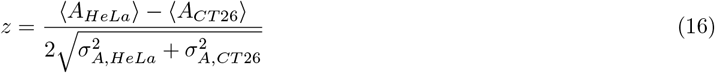

Here *σ* is the standard deviation SD.

#### M 3.4 Correlation analysis

Correlation analysis was performed using the corrcoef function of the python package numpy.^58^

#### M 3.5 Principal component analysis PCA

In order to see if the complexity of the 6-parameter fingerprint log *E, γ*, log *A, α*, log *B, β* can be further reduced, we performed a principal component analysis on the data. To this end, we utilized the function `sklearn.decomposition.PCA` from the Python module scikit-learn.^59^ The data was normalized prior to the PCA using the `sklearn.preprocessing.StandardSca` method.

### M 4 Cell culture and bead insertion

A549, C2C12, CT26, HeLa, and MDCK cells were cultured in Dulbecco’s modified Eagle medium (DMEM, Capricorn) supplemented with 1% Penicillin Streptomycin (Gibco) and 10% fetal bovine serum (FBS, Sigma-Aldrich) at a temperature of 37°C and 5% CO_2_.

HoxB8 cells, provided by the Institute for Immunology at the University of Münster, Germany, were cultured according to a protocol described elsewhere.^11^ To induce differentiation into macrophages, HoxB8 cells were suspended in differentiation medium three days prior to the experiment. After this, the same procedure as for the other cell lines, with the exception of using EDTA instead of Trypsin, was applied.

Cells were split when they reached near full confluence and seeded onto Fibronectin-coated glass cover slips (22×55×0.15 mm VWR). Medium was then aspirated, and a fresh 1:10,000 dilution of 1 µm carboxylated beads (FCDG006, Bangslabs) in medium was added to the sample. Cells were incubated for up to 15 hours to allow for adequate bead phagocytosis. Then, cells were washed with PBS to remove any residual extracellular beads. Following aspiration of PBS, a cover slip was fixed to the estimated cell spreading area using two layers of 200 µm adhesive tape (DST1950, Thorlabs, New Jersey, USA).

The sample chamber was filled with CO_2_-independent medium (CO_2_ Independent Medium, Gibco™) to ensure optimal cell viability during the experiments.

### M 5 Pharmacological treatment

Cytochalasin B (Sigma-Aldrich), Latrunculin A (Sigma-Aldrich), and Nocodazole (Sigma-Aldrich) were used to perturb the cytoskeleton. Cytochalasin B inhibits actin polymerization by capping filament ends, whereas Latrunculin A binds to actin monomers and thereby induces actin depolymerization. Nocodazole disrupts the microtubule network by binding to *β*-tubulin and inhibiting microtubule polymerization.

Cells were subjected to one of five pharmacological treatments: Cytochalasin B (CytoB 10 µg mL^−1^), Latrunculin A (LatA 1*µ*M), Nocodazole (Noc 10 µg mL^−1^), Cytochalasin B + Nocodazole (CytoB+Noc), or Latrunculin A + Nocodazole (LatA+Noc). All treatments were applied for 60 min prior to the measurements. Combined treatments were performed using the same drug concentrations as in the respective single-drug conditions.

### M 6 Immunofluorescence staining and imaging

To validate the cytoskeletal perturbations, cells were fixed and subjected to immunofluorescence staining for F-actin, microtubules, and nuclei. Following drug treatment, cells were washed three times with PBS and fixed with 4% paraformaldehyde (PFA) for 15 min at room temperature. Samples were subsequently washed with PBS and permeabilized in PBS containing 10% goat serum and 0.2% Triton X-100 for 10 min. Cells were then incubated in a blocking solution consisting of 10% goat serum and 0.2% Tween-20 in PBS.

Microtubules were stained using a mouse anti-*α*-tubulin primary antibody (A11126, Invitrogen; 1:100 dilution) for 2 h at room temperature, followed by incubation with an Alexa Fluor 568-conjugated goat anti-mouse secondary antibody (A21124, Invitrogen; 1:500 dilution) for 90 min. F-actin was labeled using Phalloidin-iFluor 647 (ab176759, Abcam; 1:500 dilution), and nuclei were stained with Hoechst (H1399, Thermo Fisher Scientific; 1:2500 dilution). All staining steps were performed protected from light. After staining, samples were washed three times with PBS and stored at 4^°^C until imaging.

Fluorescence image stacks were acquired using a spinning-disk confocal microscope (CSU-W1, Yokogawa) mounted on a Nikon Ti-1 microscope equipped with a 60× WI objective. Z-stacks were recorded with a spacing of 270 nm between adjacent planes.

## Acknowledgements

We thank the Institute for Immunology at the University of Münster for providing us with HoxB8 cells and the corre-sponding resources. T.M.M., B.E.V. and T.B. have received funding from the European Research Council (ERC) under the European Union’s Horizon 2020 research and innovation program (PolarizeMe, Grant agreement No. 771201). T.M.M., B.E.V. and T.B. have received funding from the Deutsche Forschungsgemeinschaft (DFG, German Research Foundation) – Project number 516046415 and 449750155 – RTG 2756, Projects B.3.

## Author information

### Contributions

T.M.M. conceived and built the experimental setup, performed the experiments and data analysis, interpreted the results, and drafted the manuscript.

K.v.R. contributed to cell culture, optimized the immunostaining protocols, and performed the immunostaining experiments.

B.E.V. contributed to the design and construction of the experimental setup, interpreted the results, and contributed to manuscript preparation.

T.B. conceived the study, interpreted the results, and contributed to manuscript preparation. All authors discussed the results, revised the manuscript, and approved the final version.

### M 7 Ethics declaration

### M 8 Competing interests

No competing interests

## Appendix 1 Appendix Text

### Appendix 1.1 Accounting for calibration error during active measurements

As described in Section M 2, the position of probe particles was determined using a weak detection laser in combination with back-focal-plane interferometry. The voltage signal generated by the quadrant photodiode (QPD) is proportional to the displacement of the particle relative to the center of the detection laser. To determine the proportionality constant between voltage and particle displacement, denoted by Ξ, the setup was calibrated before each experiment by moving the sample stage over a known distance while recording the QPD signal. The resulting calibration curve was then used to convert voltage signals into particle displacements. In addition, the calibration was used to reposition the tracer particle between subsequent active and passive rheology measurements.

During the course of an experiment, however, active particle motion and small positional changes relative to the detection laser can introduce systematic errors in the effective calibration factor Ξ. To account for such variations, we applied a two-step correction procedure.

First, we exploited the fact that the high-frequency part of the power spectral density (PSD) should remain unchanged between consecutive recordings of the same particle. The calibration factor Ξ of each recording was therefore adjusted such that the high-frequency PSD overlapped with that of the first measurement, which was performed immediately after the initial calibration.

Second, we utilized the fact that molecular motors operate on finite timescales and are therefore not expected to contribute significantly to intracellular activity at frequencies above approximately 500 Hz.^39^ In this frequency regime, the fluctuation-dissipation theorem (FDT) is expected to hold. We therefore applied a global rescaling of the calibration factors used in both active and passive measurements such that the average effective energy converged to the thermal limit and the FDT was fulfilled in the frequency range between 512 Hz and 1024 Hz.

## Appendix 2 Supplementary Tables

### Appendix 2.1 Principal component analysis

**Appendix Table 1:** Results of principal component analysis. The explained variance ratio quantifies how much infor-mation is captured by the corresponding component. The composition of each component is explained by its axis in parameter space. Here, the values are shown to the third decimal digit.

| Component | Explained variance ratio | $\log E$ | Normalized principal axis in parameter space | | | | |
| --- | --- | --- | --- | --- | --- | --- | --- |
| | | | $-\gamma$ | $\log A$ | $\alpha$ | $\log B$ | $\beta$ |
| PC1 | 0.515000 | -0.132000 | 0.071000 | 0.086000 | -0.223000 | 0.244000 | -0.244000 |
| PC2 | 0.269000 | -0.303000 | 0.319000 | -0.173000 | 0.109000 | -0.047000 | 0.049000 |
| PC3 | 0.146000 | 0.017000 | 0.241000 | 0.500000 | 0.048000 | -0.074000 | 0.120000 |
| PC4 | 0.041000 | -0.243000 | -0.235000 | 0.121000 | 0.265000 | 0.083000 | -0.053000 |
| PC5 | 0.028000 | 0.220000 | 0.146000 | -0.042000 | 0.261000 | 0.201000 | -0.130000 |
| PC6 | 0.001000 | -0.020000 | -0.024000 | -0.031000 | -0.062000 | 0.407000 | 0.457000 |

### Appendix 2.2 Statistics on rheology experiments

**Appendix Table 2:** Statistics on rheology experiments.

| Celltype | # Measurements | # Experiments |
| --- | --- | --- |
| HeLa | 76 | 4 |
| Cytochalasin B | 59 | 3 |
| Latrunculin A | 56 | 4 |
| Nocodazole | 59 | 3 |
| CytoB + Noc | 60 | 3 |
| LatA + Noc | 60 | 3 |
| A549 | 60 | 3 |
| C2C12 | 58 | 3 |
| CT26 | 59 | 3 |
| HoxB8 | 59 | 4 |
| MCF7 | 59 | 3 |
| MDCK | 62 | 3 |

### Appendix 2.3 Significance tests

**Appendix Table 3:**
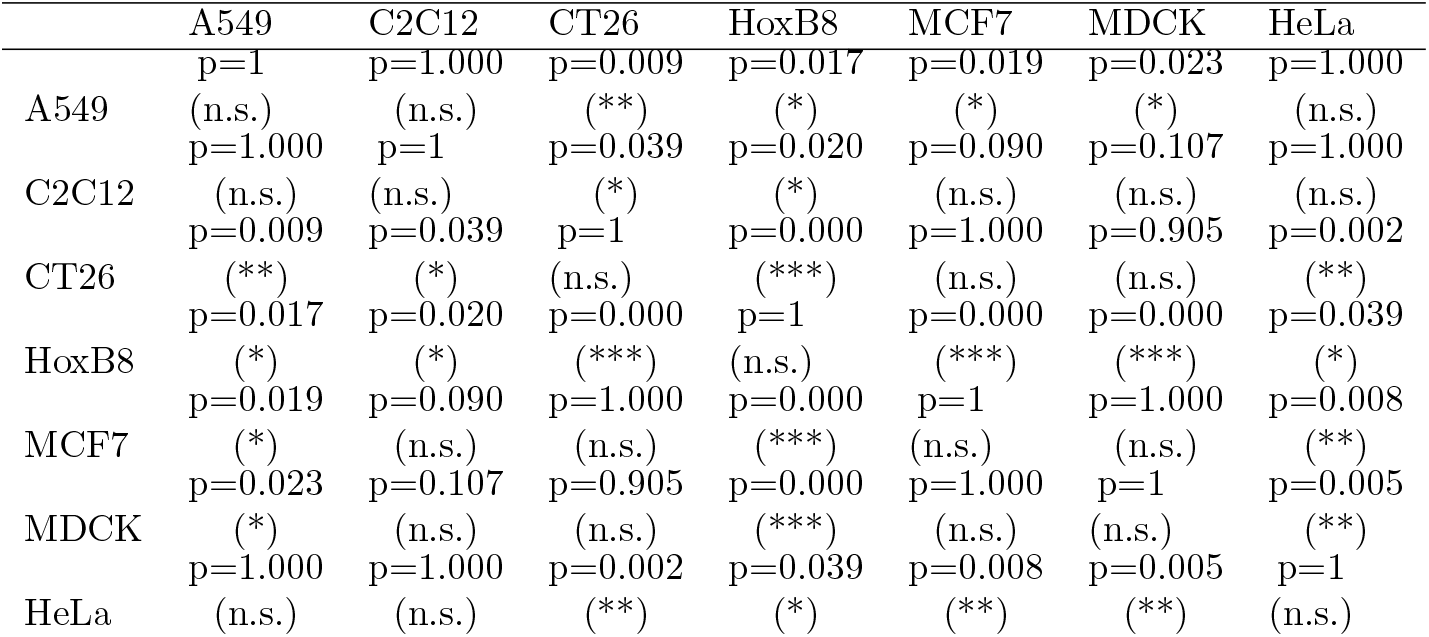
Parameter *E*: different cell types.

**Appendix Table 4:** Parameter − *γ*: different cell types.

|  | A549 | C2C12 | CT26 | HoxB8 | MCF7 | MDCK | HeLa |
| --- | --- | --- | --- | --- | --- | --- | --- |
|  | p=1 | p=1.000 | p=1.000 | p=0.002 | p=0.214 | p=1.000 | p=0.655 |
| A549 | (n.s.) | (n.s.) | (n.s.) | (**) | (n.s.) | (n.s.) | (n.s.) |
|  | p=1.000 | p=1 | p=1.000 | p=0.004 | p=0.008 | p=1.000 | p=1.000 |
| C2C12 | (n.s.) | (n.s.) | (n.s.) | (**) | (**) | (n.s.) | (n.s.) |
|  | p=1.000 | p=1.000 | p=1 | p=0.002 | p=0.350 | p=1.000 | p=0.855 |
| CT26 | (n.s.) | (n.s.) | (n.s.) | (**) | (n.s.) | (n.s.) | (n.s.) |
|  | p=0.002 | p=0.004 | p=0.002 | p=1 | p=0.000 | p=0.027 | p=0.052 |
| HoxB8 | (**) | (**) | (**) | (n.s.) | (***) | (*) | (n.s.) |
|  | p=0.214 | p=0.008 | p=0.350 | p=0.000 | p=1 | p=0.031 | p=0.008 |
| MCF7 | (n.s.) | (**) | (n.s.) | (***) | (n.s.) | (*) | (**) |
|  | p=1.000 | p=1.000 | p=1.000 | p=0.027 | p=0.031 | p=1 | p=1.000 |
| MDCK | (n.s.) | (n.s.) | (n.s.) | (*) | (*) | (n.s.) | (n.s.) |
|  | p=0.655 | p=1.000 | p=0.855 | p=0.052 | p=0.008 | p=1.000 | p=1 |
| HeLa | (n.s.) | (n.s.) | (n.s.) | (n.s.) | (**) | (n.s.) | (n.s.) |

**Appendix Table 5:** Parameter *A*: different cell types.

|  | A549 | C2C12 | CT26 | HoxB8 | MCF7 | MDCK | HeLa |
| --- | --- | --- | --- | --- | --- | --- | --- |
|  | p=1 | p=0.658 | p=0.377 | p=0.003 | p=0.000 | p=0.027 | p=0.432 |
| A549 | (n.s.) | (n.s.) | (n.s.) | (**) | (***) | (*) | (n.s.) |
|  | p=0.658 | p=1 | p=0.018 | p=0.000 | p=0.000 | p=0.000 | p=0.043 |
| C2C12 | (n.s.) | (n.s.) | (*) | (***) | (***) | (***) | (*) |
|  | p=0.377 | p=0.018 | p=1 | p=1.000 | p=1.000 | p=1.000 | p=1.000 |
| CT26 | (n.s.) | (*) | (n.s.) | (n.s.) | (n.s.) | (n.s.) | (n.s.) |
|  | p=0.003 | p=0.000 | p=1.000 | p=1 | p=1.000 | p=1.000 | p=1.000 |
| HoxB8 | (**) | (***) | (n.s.) | (n.s.) | (n.s.) | (n.s.) | (n.s.) |
|  | p=0.000 | p=0.000 | p=1.000 | p=1.000 | p=1 | p=1.000 | p=1.000 |
| MCF7 | (***) | (***) | (n.s.) | (n.s.) | (n.s.) | (n.s.) | (n.s.) |
|  | p=0.027 | p=0.000 | p=1.000 | p=1.000 | p=1.000 | p=1 | p=1.000 |
| MDCK | (*) | (***) | (n.s.) | (n.s.) | (n.s.) | (n.s.) | (n.s.) |
|  | p=0.432 | p=0.043 | p=1.000 | p=1.000 | p=1.000 | p=1.000 | p=1 |
| HeLa | (n.s.) | (*) | (n.s.) | (n.s.) | (n.s.) | (n.s.) | (n.s.) |

**Appendix Table 6:**
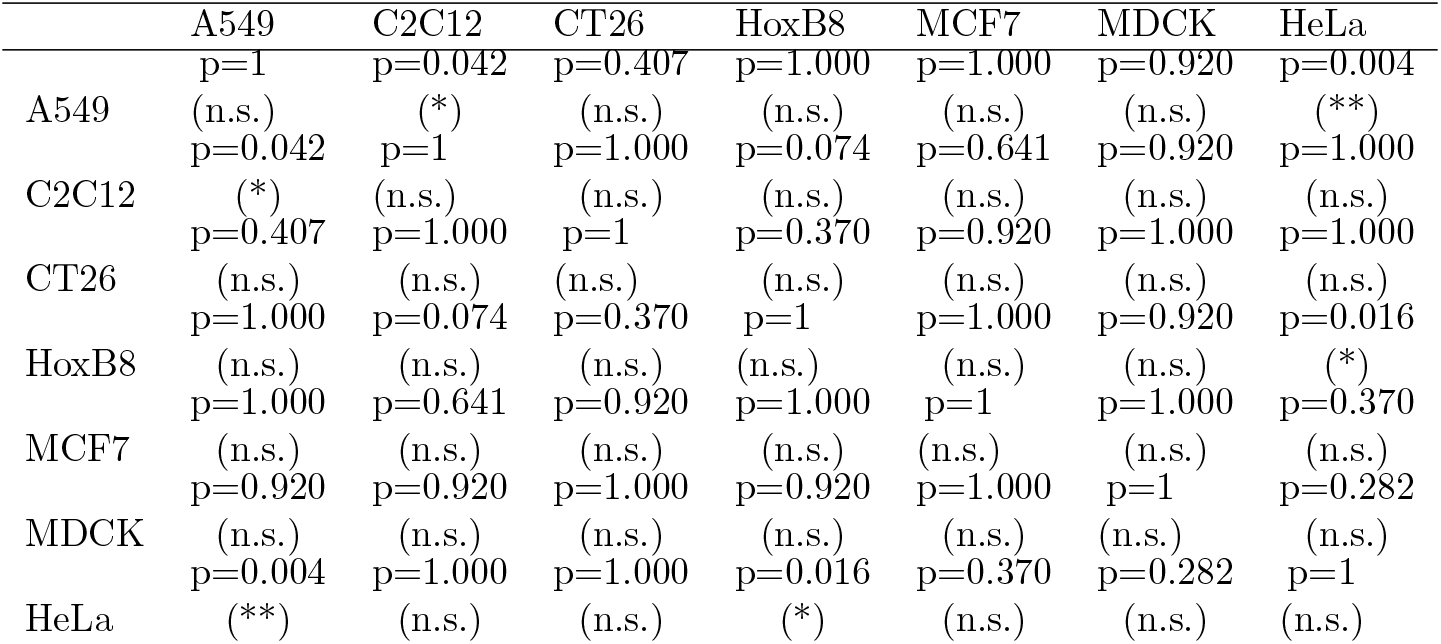
Parameter *α*: different cell types.

**Appendix Table 7:** Parameter *B*: different cell types.

|  | A549 | C2C12 | CT26 | HoxB8 | MCF7 | MDCK | HeLa |
| --- | --- | --- | --- | --- | --- | --- | --- |
|  | p=1 | p=0.265 | p=0.434 | p=1.000 | p=1.000 | p=0.904 | p=1.000 |
| A549 | (n.s.) | (n.s.) | (n.s.) | (n.s.) | (n.s.) | (n.s.) | (n.s.) |
|  | p=0.265 | p=1 | p=1.000 | p=0.739 | p=1.000 | p=1.000 | p=1.000 |
| C2C12 | (n.s.) | (n.s.) | (n.s.) | (n.s.) | (n.s.) | (n.s.) | (n.s.) |
|  | p=0.434 | p=1.000 | p=1 | p=0.904 | p=1.000 | p=1.000 | p=1.000 |
| CT26 | (n.s.) | (n.s.) | (n.s.) | (n.s.) | (n.s.) | (n.s.) | (n.s.) |
|  | p=1.000 | p=0.739 | p=0.904 | p=1 | p=1.000 | p=1.000 | p=1.000 |
| HoxB8 | (n.s.) | (n.s.) | (n.s.) | (n.s.) | (n.s.) | (n.s.) | (n.s.) |
|  | p=1.000 | p=1.000 | p=1.000 | p=1.000 | p=1 | p=1.000 | p=1.000 |
| MCF7 | (n.s.) | (n.s.) | (n.s.) | (n.s.) | (n.s.) | (n.s.) | (n.s.) |
|  | p=0.904 | p=1.000 | p=1.000 | p=1.000 | p=1.000 | p=1 | p=1.000 |
| MDCK | (n.s.) | (n.s.) | (n.s.) | (n.s.) | (n.s.) | (n.s.) | (n.s.) |
|  | p=1.000 | p=1.000 | p=1.000 | p=1.000 | p=1.000 | p=1.000 | p=1 |
| HeLa | (n.s.) | (n.s.) | (n.s.) | (n.s.) | (n.s.) | (n.s.) | (n.s.) |

**Appendix Table 8:** Parameter *β*: different cell types.

|  | A549 | C2C12 | CT26 | HoxB8 | MCF7 | MDCK | HeLa |
| --- | --- | --- | --- | --- | --- | --- | --- |
|  | p=1 | p=0.105 | p=0.126 | p=1.000 | p=1.000 | p=0.473 | p=0.473 |
| A549 | (n.s.) | (n.s.) | (n.s.) | (n.s.) | (n.s.) | (n.s.) | (n.s.) |
|  | p=0.105 | p=1 | p=1.000 | p=0.491 | p=1.000 | p=1.000 | p=1.000 |
| C2C12 | (n.s.) | (n.s.) | (n.s.) | (n.s.) | (n.s.) | (n.s.) | (n.s.) |
|  | p=0.126 | p=1.000 | p=1 | p=0.473 | p=0.916 | p=1.000 | p=1.000 |
| CT26 | (n.s.) | (n.s.) | (n.s.) | (n.s.) | (n.s.) | (n.s.) | (n.s.) |
|  | p=1.000 | p=0.491 | p=0.473 | p=1 | p=1.000 | p=1.000 | p=1.000 |
| HoxB8 | (n.s.) | (n.s.) | (n.s.) | (n.s.) | (n.s.) | (n.s.) | (n.s.) |
|  | p=1.000 | p=1.000 | p=0.916 | p=1.000 | p=1 | p=1.000 | p=1.000 |
| MCF7 | (n.s.) | (n.s.) | (n.s.) | (n.s.) | (n.s.) | (n.s.) | (n.s.) |
|  | p=0.473 | p=1.000 | p=1.000 | p=1.000 | p=1.000 | p=1 | p=1.000 |
| MDCK | (n.s.) | (n.s.) | (n.s.) | (n.s.) | (n.s.) | (n.s.) | (n.s.) |
|  | p=0.473 | p=1.000 | p=1.000 | p=1.000 | p=1.000 | p=1.000 | p=1 |
| HeLa | (n.s.) | (n.s.) | (n.s.) | (n.s.) | (n.s.) | (n.s.) | (n.s.) |

**Appendix Table 9:**
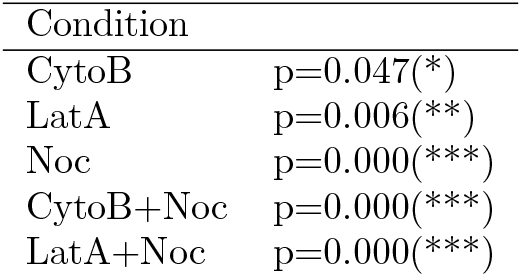
Parameter *E*: drug conditions vs HeLa.

**Appendix Table 10:** Parameter *γ*: drug conditions vs HeLa.

| Condition |  |
| --- | --- |
| CytoB | p=0.000(***) |
| LatA | p=0.000(***) |
| Noc | p=0.156(n.s.) |
| CytoB+Noc | p=0.000(***) |
| LatA+Noc | p=0.064(n.s.) |

**Appendix Table 11:** Parameter *A*: drug conditions vs HeLa.

| Condition |  |
| --- | --- |
| CytoB | p=0.085(n.s.) |
| LatA | p=0.000(***) |
| Noc | p=0.001(**) |
| CytoB+Noc | p=0.000(***) |
| LatA+Noc | p=0.000(***) |

**Appendix Table 12:**
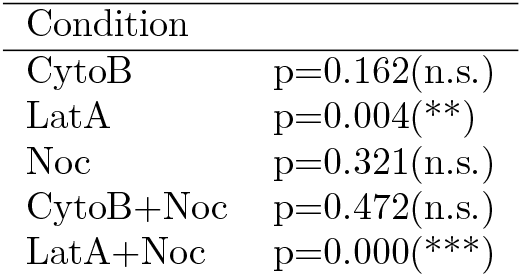
Parameter *α*: drug conditions vs HeLa.

| Condition |  |
| --- | --- |
| CytoB | p=0.162(n.s.) |
| LatA | p=0.004(**) |
| Noc | p=0.321(n.s.) |
| CytoB+Noc | p=0.472(n.s.) |
| LatA+Noc | p=0.000(***) |

**Appendix Table 13:**
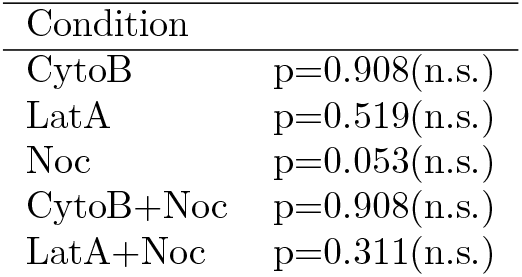
Parameter *B*: drug conditions vs HeLa.

**Appendix Table 14:** Parameter *β*: drug conditions vs HeLa.

| Condition |  |
| --- | --- |
| CytoB | p=0.450(n.s.) |
| LatA | p=0.450(n.s.) |
| Noc | p=0.396(n.s.) |
| CytoB+Noc | p=0.323(n.s.) |
| LatA+Noc | p=0.450(n.s.) |

### Appendix 2.4 *R*^2^-values for fits of bootstrapped data

**Appendix Table 15:** Mean R2 values for fit quality: bootstrapped data.

| Group | $R^2$ $E_{eff}$ | $R^2$ G real | $R^2$ G imag |
| --- | --- | --- | --- |
| A549 | 0.985 | 0.997 | 0.995 |
| C2C12 | 0.996 | 0.997 | 0.996 |
| CT26 | 0.991 | 0.987 | 0.996 |
| HoxB8 | 0.987 | 0.996 | 0.995 |
| MCF7 | 0.990 | 0.993 | 0.997 |
| MDCK | 0.983 | 0.988 | 0.999 |
| HeLa | 0.966 | 0.990 | 0.998 |
| CytoB | 0.908 | 0.978 | 0.991 |
| LatA | 0.934 | 0.994 | 0.998 |
| Noc | 0.398 | 0.995 | 0.999 |
| CytoB+Noc | 0.740 | 0.973 | 0.998 |
| LatA+Noc | -0.079 | 0.951 | 0.989 |

### Appendix 2.5 *R*^2^-values for fits of not bootstrapped data

**Appendix Table 16:** Mean R2 values for fit quality: raw data.

| Group | $R^2$ $E_{eff}$ | $R^2$ G real | $R^2$ G imag |
| --- | --- | --- | --- |
| A549 | 0.882 | 0.949 | 0.980 |
| C2C12 | 0.938 | 0.965 | 0.985 |
| CT26 | 0.910 | 0.961 | 0.981 |
| HoxB8 | 0.918 | 0.958 | 0.977 |
| MCF7 | 0.912 | 0.960 | 0.985 |
| MDCK | 0.926 | 0.951 | 0.983 |
| HeLa | 0.942 | 0.958 | 0.982 |
| CytoB | 0.656 | 0.980 | 0.993 |
| LatA | 0.822 | 0.976 | 0.991 |
| Noc | 0.212 | 0.980 | 0.993 |
| CytoB+Noc | 0.367 | 0.989 | 0.997 |
| LatA+Noc | 0.114 | 0.991 | 0.997 |

**Appendix Table 17:** Biological characteristics of the investigated cell lines.

| Celltype | Species | Cell Type | Cancer Level | Metastatic Potential | Migratory Potential |
| --- | --- | --- | --- | --- | --- |
| A549 | Human | Epithelial | Cancer | Moderate | Low |
| C2C12 | Mouse | Mesenchymal | Non-cancer | Low | Moderate |
| CT26 | Mouse | Intermediate | Cancer | High | Moderate |
| HoxB8 | Mouse | Neither | Non-cancer | Low | High |
| MCF7 | Human | Epithelial | Cancer | Low | Low |
| MDCK | Dog | Epithelial | Non-cancer | Low | Low |
| HeLa | Human | Intermediate | Cancer | Moderate | Moderate |

## Appendix 3 Figures

**Appendix Figure 1:**
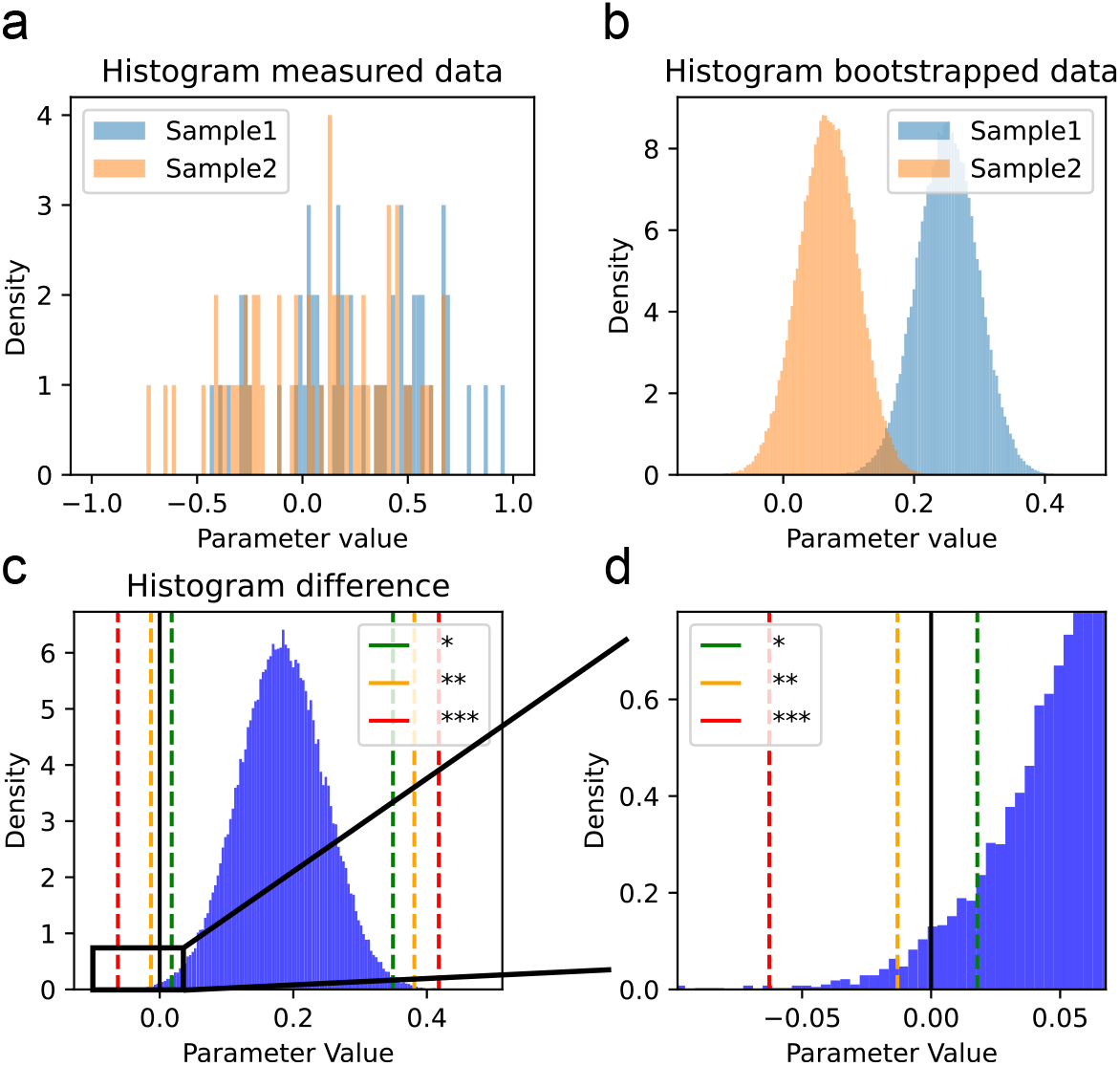
Schematic of bootstrapping significance test. a) Histogram for the estimates of two different parameters. b) Bootstrapping procedure was performed on both parameters to get an estimate for the distribution of the mean. c) Difference of both distribution is calculated. d) Depending on which percentile of the distribution is below 0, the significance score is determined.

**Appendix Figure 2:**
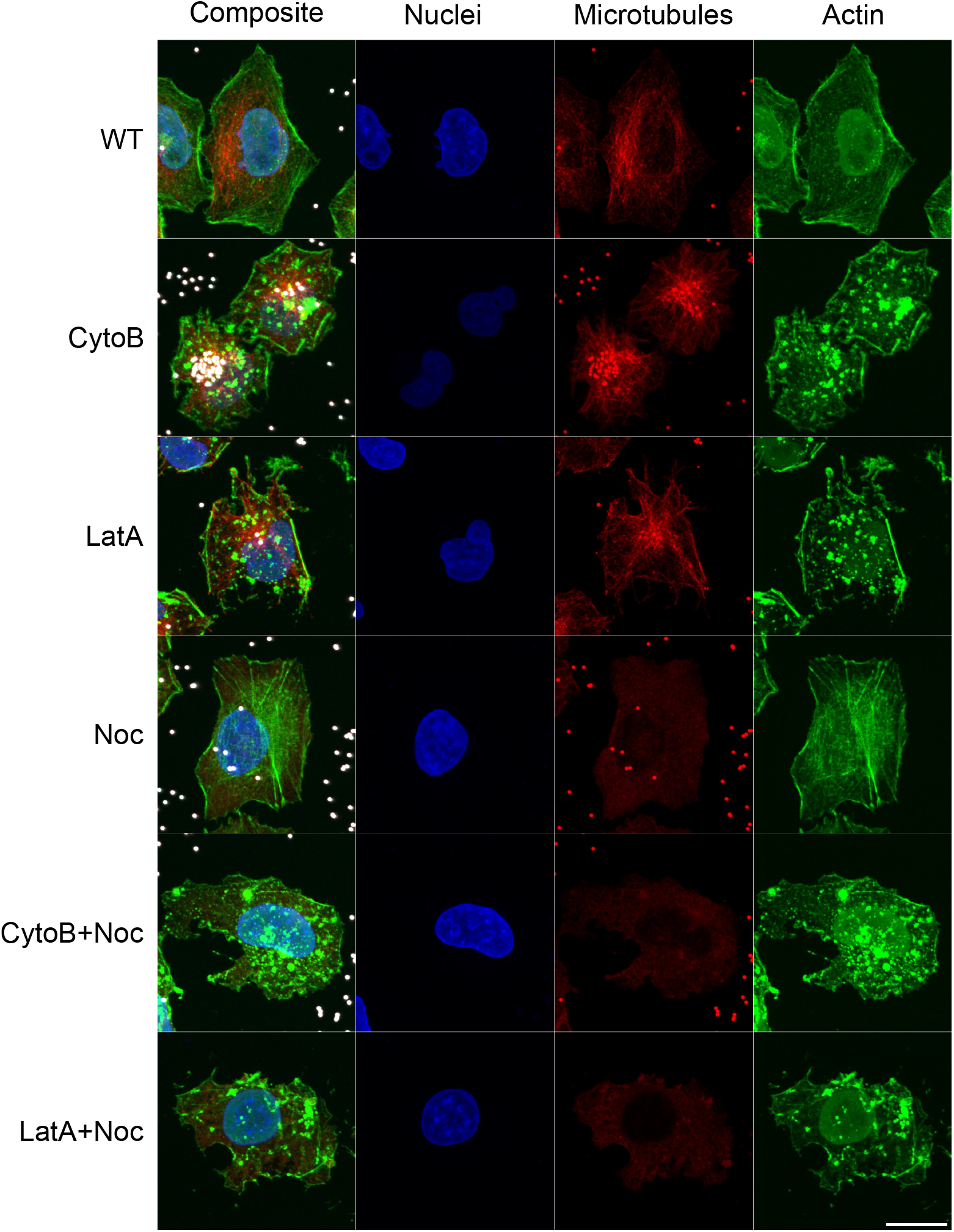
Representative images of immunostainings on HeLa WT cells treated with 10 *µ*g*/*mL Cytochalasin B (CytoB), 1*µ*M Latrunculin A (LatA), 10 *µ*g*/*mL Nocodazole (Noc) as well as the combinations CytoB+Noc and LatA+Noc. The composite images show the overlay of DNA labeled using Hoechst (blue), microtubules stained with anti-*α*-tubulin primary antibody with Alexa Fluor 568 (red), actin stained with Phalloidin 647 (green), and 1 *µ*m carboxylated beads 480 nm (white). The wide emission spectrum of the beads causes them to appear in the microtubule channel. Scale bar: 20 µm.

**Appendix Figure 3:**
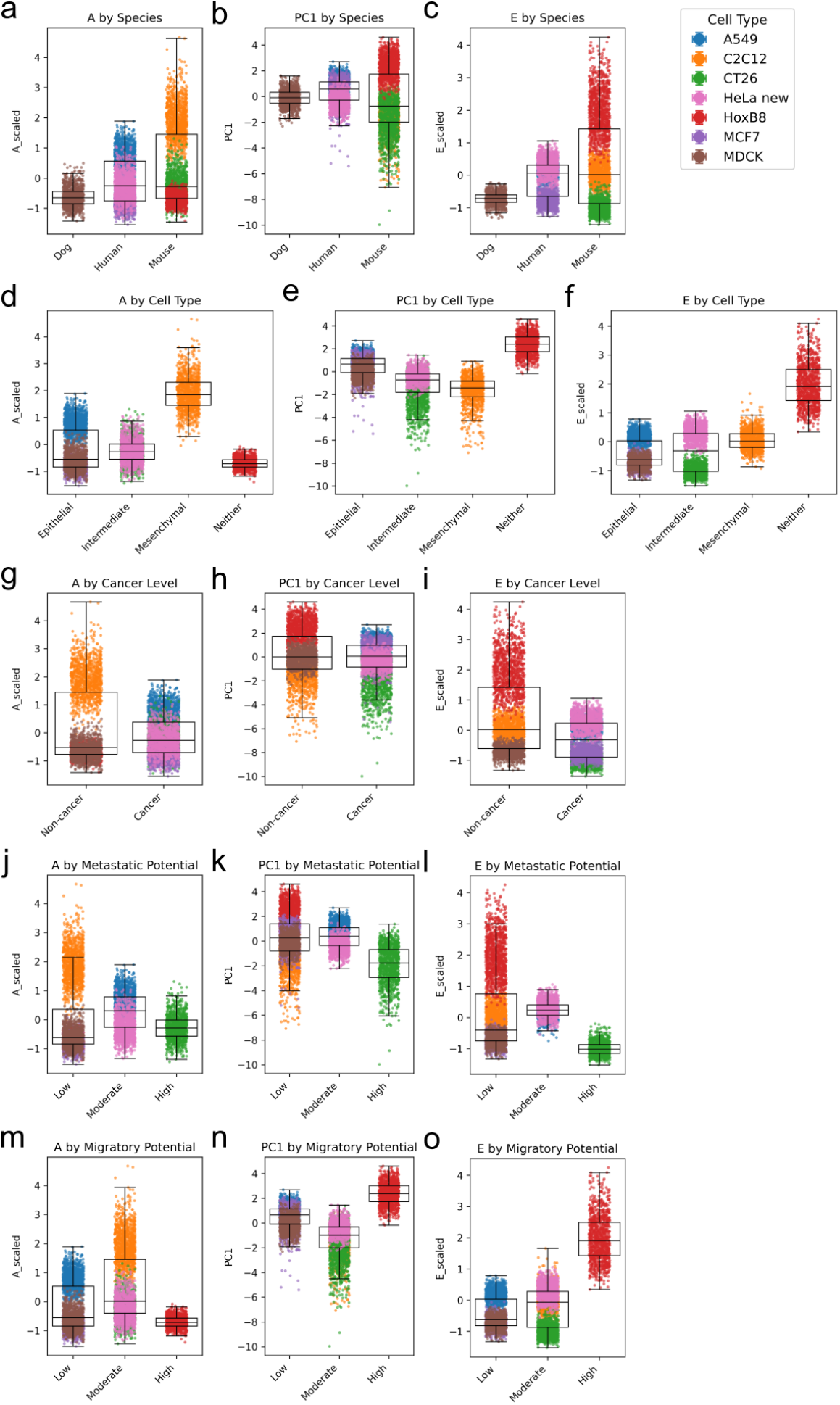
Distribution of the phase-space parameters resistance (log *A*), fluidity (PC1), and activity (log *E*) after grouping the investigated cell types according to a-c) species, d-f) cell type, g-i) cancer status, j-l) metastatic potential, and m-o) migratory potential.

## Notes

### Competing Interest Statement

The authors have declared no competing interest.

### Summary of Updates

Repeated measurements on wild-type HeLa cells. Repeated all Cytochalasin B and Nocodazole experiments and increased the number of analyzed cells to approximately 60 per condition. Performed additional experiments using Latrunculin A and combined Latrunculin A + Nocodazole treatment. Performed immunostainings for all cytoskeletal perturbation conditions (WT, Cytochalasin B, Latrunculin A, Nocodazole, Cytochalasin B + Nocodazole, and Latrunculin A + Nocodazole). Refined the rheological analysis procedure and expanded the methodological description. Revised the manuscript text throughout and expanded the discussion of limitations and biological interpretation.

## References

[1] C.-P. Heisenberg and Y. Bellaïche, “Forces in tissue morphogenesis and patterning,” Cell, vol. 153, pp. 948–962, May 2013.

[2] L. G. Pimpale, T. C. Middelkoop, A. Mietke, and S. W. Grill, “Cell lineage-dependent chiral actomyosin flows drive cellular rearrangements in early caenorhabditis elegans development,” eLife, vol. 9, July 2020.

[3] G. G. Gundersen and H. J. Worman, “Nuclear positioning,” Cell, vol. 152, pp. 1376–1389, Mar. 2013.

[4] S. A. Ruiz and C. S. Chen, “Emergence of patterned stem cell differentiation within multicellular structures,” Stem Cells, vol. 26, pp. 2921–2927, Nov. 2008.

[5] D. T. Tambe, C. C. Hardin, T. E. Angelini, K. Rajendran, C. Y. Park, X. Serra-Picamal, E. H. Zhou, M. H. Zaman, J. P. Butler, D. A. Weitz, J. J. Fredberg, and X. Trepat, “Collective cell guidance by cooperative intercellular forces,” Nat. Mater., vol. 10, pp. 469–475, June 2011.

[6] A. D. Doyle, F. W. Wang, K. Matsumoto, and K. M. Yamada, “One-dimensional topography underlies three-dimensional fibrillar cell migration,” Journal of Cell Biology, vol. 184, pp. 481–490, Feb 2009.

[7] A. Pitaval, Q. Tseng, M. Bornens, and M. Théry, “Cell shape and contractility regulate ciliogenesis in cell cycle-arrested cells,” J. Cell Biol., vol. 191, pp. 303–312, Oct. 2010.

[8] G. Fläschner, C. I. Roman, N. Strohmeyer, D. Martinez-Martin, and D. J. Müller, “Rheology of rounded mammalian cells over continuous high-frequencies,” Nature Communications, vol. 12, p. 2922, May 2021.

[9] M. Li, D. Dang, L. Liu, N. Xi, and Y. Wang, “Atomic force microscopy in characterizing cell mechanics for biomedical applications: A review,” IEEE Transactions on NanoBioscience, vol. 16, pp. 523–540, Sept. 2017.

[10] E. Evans and A. Yeung, “Apparent viscosity and cortical tension of blood granulocytes determined by micropipet aspiration,” Biophysical Journal, vol. 56, pp. 151–160, July 1989.

[11] H. Schürmann, F. Abbasi, A. Russo, A. D. Hofemeier, M. Brandt, J. Roth, T. Vogl, and T. Betz, “Analysis of monocyte cell tractions in 2.5d reveals mesoscale mechanics of podosomes during substrate-indenting cell protrusion,” Journal of Cell Science, vol. 135, p. jcs259042, May 2022.

[12] H. D. Belly, A. Stubb, A. Yanagida, C. Labouesse, P. H. Jones, E. K. Paluch, and K. J. Chalut, “Membrane tension gates ERK-mediated regulation of pluripotent cell fate,” Cell Stem Cell, vol. 28, pp. 273–284.e6, Feb. 2021.

[13] N. Wang, I. M. Tolić-Nørrelykke, J. Chen, S. M. Mijailovich, J. P. Butler, J. J. Fredberg, and D. Stamenović, “Cell prestress. i. stiffness and prestress are closely associated in adherent contractile cells,” American Journal of Physiology-Cell Physiology, vol. 282, pp. C606–C616, Mar 2002.

[14] P. Rosendahl, K. Plak, A. Jacobi, M. Kraeter, N. Toepfner, O. Otto, C. Herold, M. Winzi, M. Herbig, Y. Ge, S. Girardo, K. Wagner, B. Baum, and J. Guck, “Real-time fluorescence and deformability cytometry,” Nature Methods, vol. 15, pp. 355–358, May 2018.

[15] S. Mathieu and J.-B. Manneville, “Intracellular mechanics: connecting rheology and mechanotransduction,” Current Opinion in Cell Biology, vol. 56, pp. 34–44, Feb. 2019.

[16] M. Guo, A. J. Ehrlicher, M. H. Jensen, M. Renz, J. R. Moore, R. D. Goldman, J. Lippincott-Schwartz, F. C. Mackintosh, and D. A. Weitz, “Probing the stochastic, motor-driven properties of the cytoplasm using force spectrum microscopy,” Cell, vol. 158, pp. 822–832, Aug 2014.

[17] J. Xie, J. Najafi, R. L. Borgne, J.-M. Verbavatz, C. Durieu, J. Sallé, and N. Minc, “Contribution of cytoplasm viscoelastic properties to mitotic spindle positioning,” Proceedings of the National Academy of Sciences, vol. 119, Feb. 2022.

[18] G. Volpe, O. M. Maragò, H. Rubinsztein-Dunlop, G. Pesce, A. B. Stilgoe, G. Volpe, G. Tkachenko, V. G. Truong, S. N. Chormaic, F. Kalantarifard, P. Elahi, M. Käll, A. Callegari, M. I. Marqués, A. A. R. Neves, W. L. Moreira, A. Fontes, C. L. Cesar, R. Saija, A. Saidi, P. Beck, J. S. Eismann, P. Banzer, T. F. D. Fernandes, F. Pedaci, W. P. Bowen, R. Vaippully, M. Lokesh, B. Roy, G. Thalhammer-Thurner, M. Ritsch-Marte, L. P. García, A. V. Arzola, I. P. Castillo, A. Argun, T. M. Muenker, B. E. Vos, T. Betz, I. Cristiani, P. Minzioni, P. J. Reece, F. Wang, D. McGloin, J. C. Ndukaife, R. Quidant, R. P. Roberts, C. Laplane, T. Volz, R. Gordon, D. Hanstorp, J. T. Marmolejo, G. D. Bruce, K. Dholakia, T. Li, O. Brzobohatý, S. H. Simpson, P. Zemánek, F. Ritort, Y. Roichman, V. Bobkova, R. Wittkowski, C. Denz, G. V. P. Kumar, A. Foti, M. G. Donato, P. G. Gucciardi, L. Gardini, G. Bianchi, A. V. Kashchuk, M. Capitanio, L. Paterson, P. H. Jones, K. Berg-Sørensen, Y. F. Barooji, L. B. Oddershede, P. Pouladian, D. Preece, C. B. Adiels, A. C. D. Luca, A. Magazzu, D. B. Ciriza, M. A. Iati, and G. A. Swartzlander, “Roadmap for optical tweezers,” Journal of Physics: Photonics, vol. 5, p. 022501, Apr. 2023.

[19] B. H. Blehm, A. Devine, J. R. Staunton, and K. Tanner, “In vivo tissue has non-linear rheological behavior distinct from 3d biomimetic hydrogels, as determined by AMOTIV microscopy,” Biomaterials, vol. 83, pp. 66–78, Mar. 2016.

[20] S. Hurst, B. E. Vos, M. Brandt, and T. Betz, “Intracellular softening and increased viscoelastic fluidity during division,” Nature Physics, vol. 17, pp. 1270–1276, Nov 2021.

[21] C. Battle, C. P. Broedersz, N. Fakhri, V. F. Geyer, J. Howard, C. F. Schmidt, and F. C. MacKintosh, “Broken detailed balance at mesoscopic scales in active biological systems,” Science, vol. 352, pp. 604–607, Apr. 2016.

[22] F. S. Gnesotto, F. Mura, J. Gladrow, and C. P. Broedersz, “Broken detailed balance and non-equilibrium dynamics in living systems: a review,” Reports on Progress in Physics, vol. 81, p. 066601, Apr. 2018.

[23] M. Yousafzai, V. Yadav, S. Amiri, Y. Errami, S. Amiri, and M. Murrell, “Active regulation of pressure and volume defines an energetic constraint on the size of cell aggregates,” Physical Review Letters, vol. 128, Jan. 2022.

[24] X. Fang and J. Wang, “Nonequilibrium thermodynamics in cell biology: Extending equilibrium formalism to cover living systems,” Annual Review of Biophysics, vol. 49, pp. 227–246, May 2020.

[25] H. Yamaguchi and J. Condeelis, “Regulation of the actin cytoskeleton in cancer cell migration and invasion,” Biochimica et Biophysica Acta (BBA) - Molecular Cell Research, vol. 1773, pp. 642–652, May 2007.

[26] P. P. Provenzano and P. J. Keely, “Mechanical signaling through the cytoskeleton regulates cell proliferation by coordinated focal adhesion and rho GTPase signaling,” Journal of Cell Science, vol. 124, pp. 1195–1205, Apr. 2011.

[27] M. T. Abreu-Blanco, J. J. Watts, J. M. Verboon, and S. M. Parkhurst, “Cytoskeleton responses in wound repair,” Cellular and Molecular Life Sciences, vol. 69, pp. 2469–2483, Feb. 2012.

[28] A. S. Moore, S. M. Coscia, C. L. Simpson, F. E. Ortega, E. C. Wait, J. M. Heddleston, J. J. Nirschl, C. J. Obara, P. Guedes-Dias, C. A. Boecker, T.-L. Chew, J. A. Theriot, J. Lippincott-Schwartz, and E. L. F. Holzbaur, “Actin cables and comet tails organize mitochondrial networks in mitosis,” Nature, vol. 591, pp. 659–664, Mar 2021.

[29] A. A. Jord, G. Letort, S. Chanet, F.-C. Tsai, C. Antoniewski, A. Eichmuller, C. D. Silva, J.-R. Huynh, N. S. Gov, R. Voituriez, M.-É. Terret, and M.-H. Verlhac, “Cytoplasmic forces functionally reorganize nuclear condensates in oocytes,” Nature Communications, vol. 13, Aug. 2022.

[30] G. Guigas, C. Kalla, and M. Weiss, “Probing the nanoscale viscoelasticity of intracellular fluids in living cells,” Biophysical Journal, vol. 93, pp. 316–323, July 2007.

[31] A. Bonfanti, J. L. Kaplan, G. Charras, and A. Kabla, “Fractional viscoelastic models for power-law materials,” vol. 16, no. 26, pp. 6002–6020.

[32] A. Rigato, A. Miyagi, S. Scheuring, and F. Rico, “High-frequency microrheology reveals cytoskeleton dynamics in living cells,” Nature Physics, vol. 13, pp. 771–775, Aug 2017.

[33] A. Nguyen, M. Brandt, T. M. Muenker, and T. Betz, “Multi-oscillation microrheology via acoustic force spectroscopy enables frequency-dependent measurements on endothelial cells at high-throughput,” Lab on a Chip, vol. 21, no. 10, pp. 1929–1947, 2021.

[34] T. M. Muenker, G. Knotz, M. Krüger, and T. Betz, “Accessing activity and viscoelastic properties of artificial and living systems from passive measurement,” Nat. Mater., vol. 23, pp. 1283–1291, Sept. 2024.

[35] G. Knotz, T. M. Muenker, T. Betz, and M. Krüger, “Time reversal breaking of colloidal particles in cells,” The Journal of Chemical Physics, vol. 165, p. 015102, 07 2026.

[36] G. Knotz, T. M. Muenker, T. Betz, and M. Krüger, “Evaluating non-equilibrium trajectories via mean back relaxation: Dependence on length and time scales,” J. Chem. Phys., vol. 163, p. 124118, Sept. 2025.

[37] W. W. Ahmed, É. Fodor, and T. Betz, “Active cell mechanics: Measurement and theory,” Biochimica et Biophysica Acta (BBA) - Molecular Cell Research, vol. 1853, pp. 3083–3094, Nov 2015.

[38] F. C. MacKintosh and C. F. Schmidt, “Active cellular materials,” Current Opinion in Cell Biology, vol. 22, pp. 29–35, Feb 2010.

[39] W. Junge, “ATP synthase and other motor proteins,” Proc. Natl. Acad. Sci. U. S. A., vol. 96, pp. 4735–4737, Apr. 1999.

[40] H. Turlier, D. A. Fedosov, B. Audoly, T. Auth, N. S. Gov, C. Sykes, J.-F. Joanny, G. Gompper, and T. Betz, “Equilibrium physics breakdown reveals the active nature of red blood cell flickering,” Nat. Phys., vol. 12, pp. 513–519, May 2016.

[41] D. Mizuno, C. Tardin, C. F. Schmidt, and F. C. Mackintosh, “Nonequilibrium mechanics of active cytoskeletal networks,” Science, vol. 315, pp. 370–373, Jan. 2007.

[42] W. W. Ahmed, É. Fodor, M. Almonacid, M. Bussonnier, M.-H. Verlhac, N. Gov, P. Visco, F. van Wijland, and T. Betz, “Active mechanics reveal molecular-scale force kinetics in living oocytes,” Biophys. J., vol. 114, pp. 1667–1679, Apr. 2018.

[43] C. Rotsch and M. Radmacher, “Drug-induced changes of cytoskeletal structure and mechanics in fibroblasts: an atomic force microscopy study,” Biophys. J., vol. 78, pp. 520–535, Jan. 2000.

[44] H. W. Wu, T. Kuhn, and V. T. Moy, “Mechanical properties of L929 cells measured by atomic force microscopy: effects of anticytoskeletal drugs and membrane crosslinking,” Scanning, vol. 20, pp. 389–397, Aug. 1998.

[45] I. Spector, N. R. Shochet, D. Blasberger, and Y. Kashman, “Latrunculins–novel marine macrolides that disrupt microfilament organization and affect cell growth: I. comparison with cytochalasin D,” Cell Motil. Cytoskeleton, vol. 13, no. 3, pp. 127–144, 1989.

[46] M. E. Grady, R. J. Composto, and D. M. Eckmann, “Cell elasticity with altered cytoskeletal architectures across multiple cell types,” J. Mech. Behav. Biomed. Mater., vol. 61, pp. 197–207, Aug. 2016.

[47] H. Ebata, K. Umeda, K. Nishizawa, W. Nagao, S. Inokuchi, Y. Sugino, T. Miyamoto, and D. Mizuno, “Activity-dependent glassy cell mechanics i: Mechanical properties measured with active microrheology,” Biophys. J., vol. 122, pp. 1781–1793, May 2023.

[48] K. Umeda, K. Nishizawa, W. Nagao, S. Inokuchi, Y. Sugino, H. Ebata, and D. Mizuno, “Activity-dependent glassy cell mechanics II: Nonthermal fluctuations under metabolic activity,” Biophys. J., vol. 122, pp. 4395–4413, Nov. 2023.

[49] A. E. Ekpenyong, G. Whyte, K. Chalut, S. Pagliara, F. Lautenschläger, C. Fiddler, S. Paschke, U. F. Keyser, E. R. Chilvers, and J. Guck, “Viscoelastic properties of differentiating blood cells are fate- and function-dependent,” PLOS ONE, vol. 7, pp. 1–10, 09 2012.

[50] J. Rother, H. Nöding, I. Mey, and A. Janshoff, “Atomic force microscopy-based microrheology reveals significant differences in the viscoelastic response between malign and benign cell lines,” Open Biol., vol. 4, p. 140046, May 2014.

[51] H. Chen and J. Nalbantoglu, “Ring cell migration assay identifies distinct effects of extracellular matrix proteins on cancer cell migration,” BMC Res. Notes, vol. 7, p. 183, Mar. 2014.

[52] T. G. Mason and D. A. Weitz, “Optical Measurements of Frequency-Dependent Linear Viscoelastic Moduli of Complex Fluids,” Physical Review Letters, vol. 74, pp. 1250–1253, Feb. 1995. Publisher: American Physical Society.

[53] F. Gittes, B. Schnurr, P. D. Olmsted, F. C. MacKintosh, and C. F. Schmidt, “Microscopic viscoelasticity: Shear moduli of soft materials determined from thermal fluctuations,” Phys. Rev. Lett., vol. 79, pp. 3286–3289, Oct 1997.

[54] A. Farré and M. Montes-Usategui, “A force detection technique for single-beam optical traps based on direct measurement of light momentum changes,” vol. 18, no. 11, pp. 11955–11968. Publisher: Optical Society of America.

[55] F. Marsà, A. Farré, E. Martín-Badosa, and M. Montes-Usategui, “Holographic optical tweezers combined with back-focal-plane displacement detection,” Opt. Express, vol. 21, pp. 30282–30294, Dec 2013.

[56] B. Efron, R. Tibshirani, and R. J. Tibshirani, An introduction to the bootstrap. Chapman & Hall/CRC Monographs on Statistics and Applied Probability, Philadelphia, PA: Chapman & Hall/CRC, May 1994.

[57] S. Holm, “A simple sequentially rejective multiple test procedure,” Scand. J. Statist., vol. 6, no. 2, pp. 65–70, 1979.

[58] C. R. Harris, K. J. Millman, S. J. van der Walt, R. Gommers, P. Virtanen, D. Cournapeau, E. Wieser, J. Taylor, S. Berg, N. J. Smith, R. Kern, M. Picus, S. Hoyer, M. H. van Kerkwijk, M. Brett, A. Haldane, J. F. del Río, M. Wiebe, P. Peterson, P. Gérard-Marchant, K. Sheppard, T. Reddy, W. Weckesser, H. Abbasi, C. Gohlke, and T. E. Oliphant, “Array programming with NumPy,” Nature, vol. 585, pp. 357–362, Sept. 2020.

[59] F. Pedregosa, G. Varoquaux, A. Gramfort, V. Michel, B. Thirion, O. Grisel, M. Blondel, P. Prettenhofer, R. Weiss, V. Dubourg, J. Vanderplas, A. Passos, D. Cournapeau, M. Brucher, M. Perrot, and E. Duchesnay, “Scikit-learn: Machine learning in Python,” Journal of Machine Learning Research, vol. 12, pp. 2825–2830, 2011.

