## Supplementary figures and images for "Intracellular mechanical fingerprint reveals cell type-specific mechanical differences"

### Appendix Figure 1

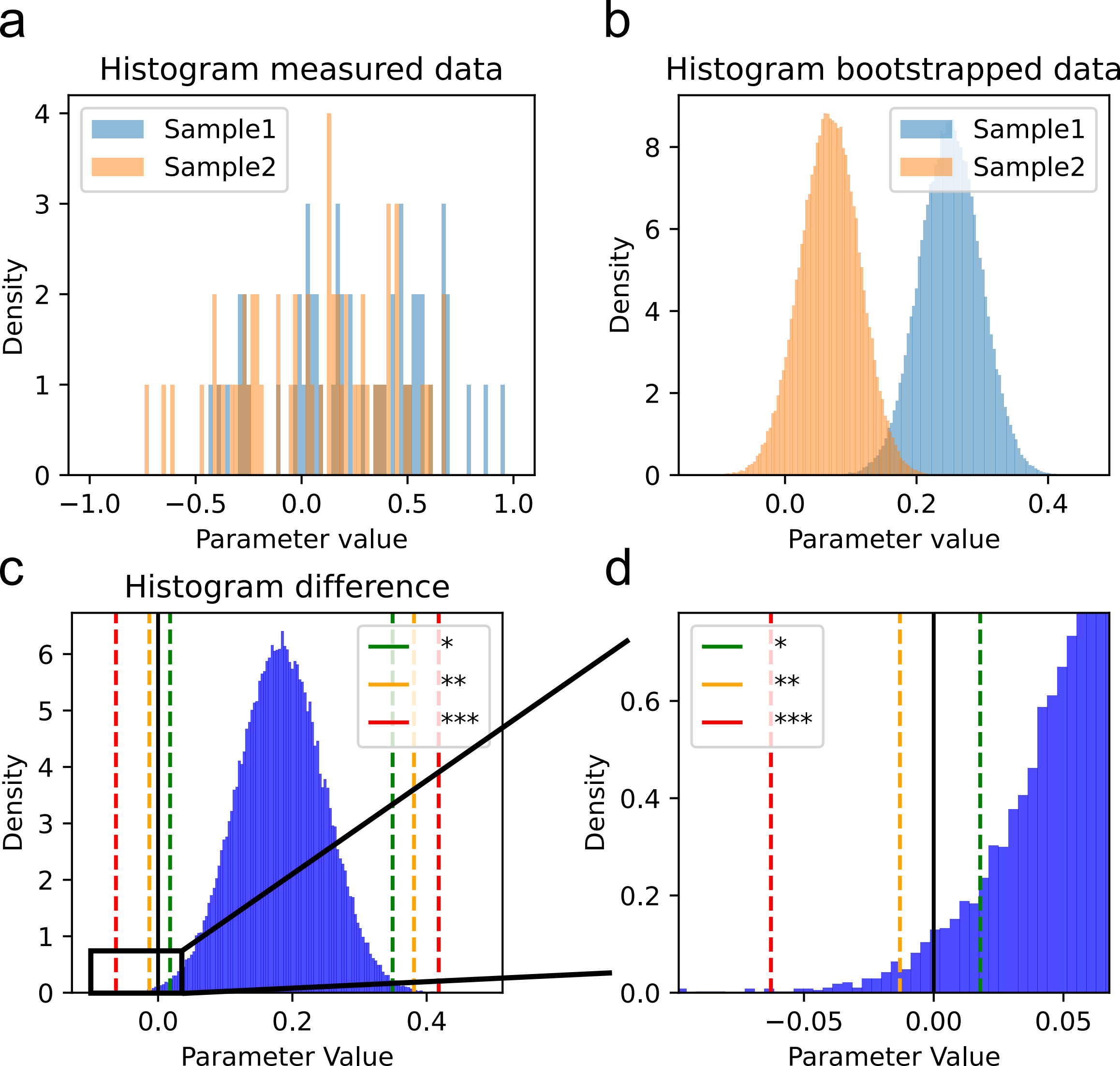

### Appendix Figure 2

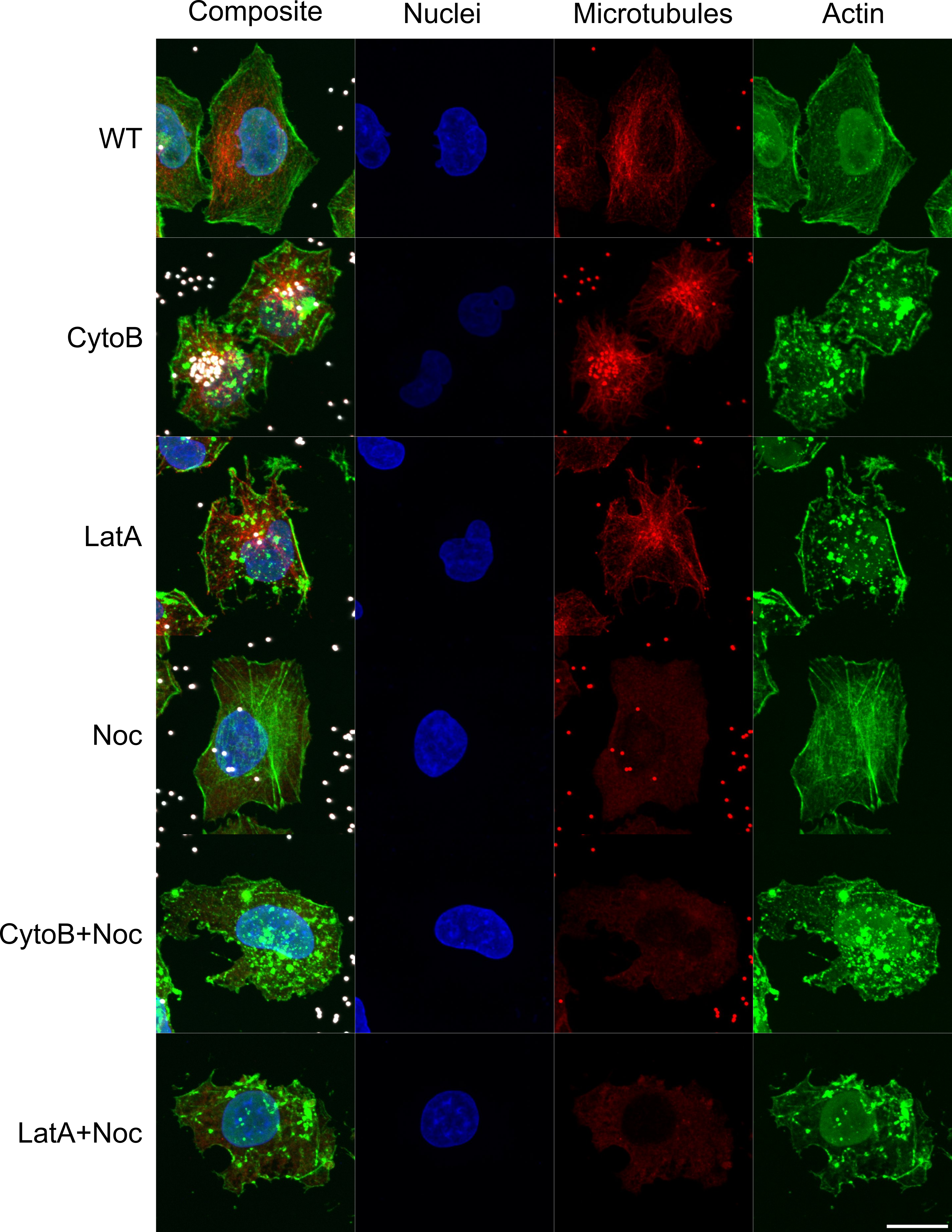

### Appendix Figure 3

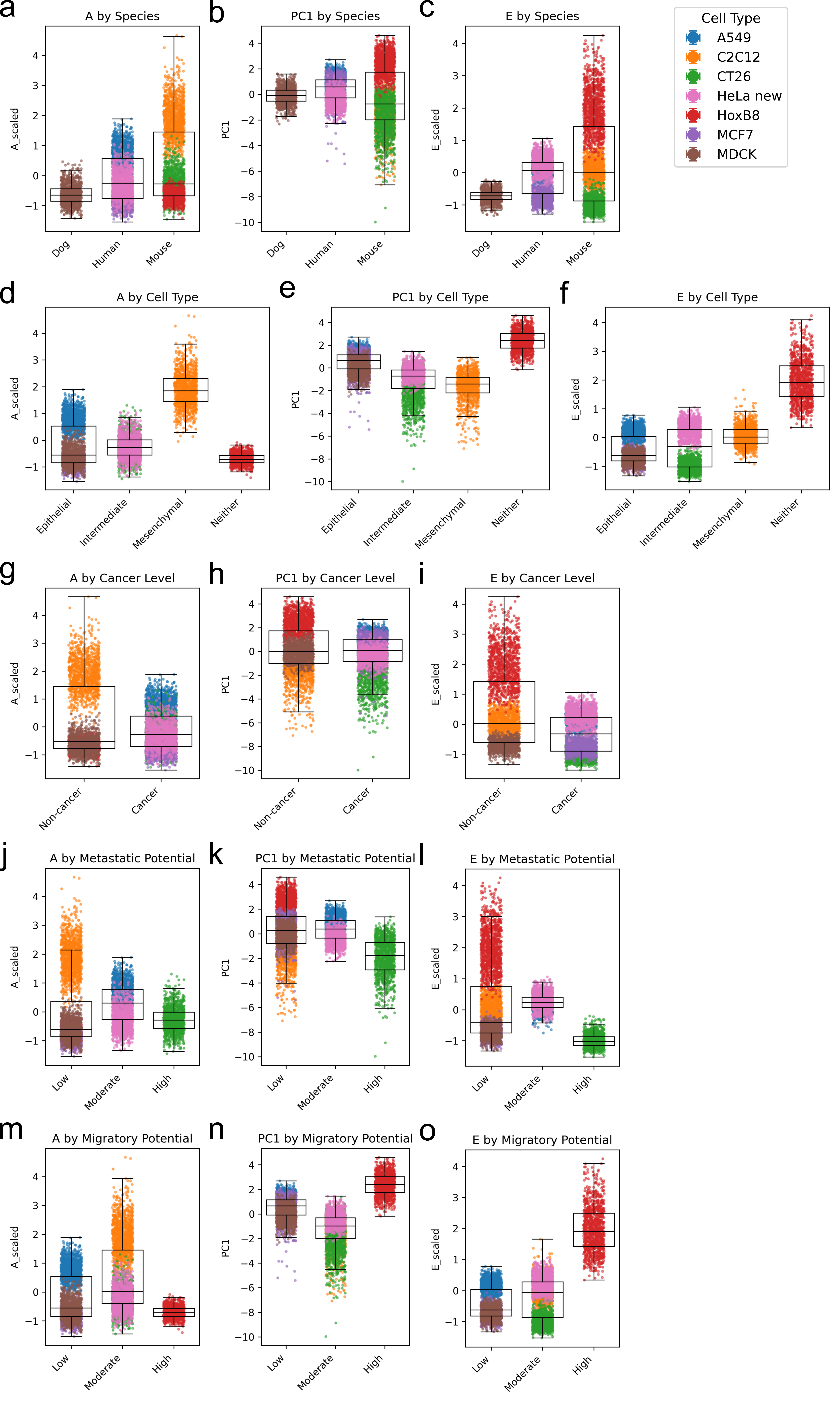
